# GroEL/ES chaperonin unfolds then encapsulates a nascent protein on the ribosome

**DOI:** 10.1101/2024.08.12.607569

**Authors:** Alžběta Roeselová, Sarah L. Maslen, Jessica Zhiyun He, Gabija Jurkeviciute, J. Mark Skehel, Radoslav Enchev, David Balchin

## Abstract

The bacterial chaperonin GroEL/ES promotes protein folding post-translation by transiently encapsulating substrate proteins within a central chamber. GroEL also binds translating ribosomes *in vivo*, suggesting an additional role in cotranslational folding. Here, we used biochemical reconstitution, structural proteomics and electron microscopy to study the mechanism by which GroEL/ES engages nascent polypeptides. We show that GroEL binds nascent chains on the inside of its cavity via the apical domains and disordered C-terminal tails, resulting in local structural destabilization of the substrate. Ribosome-tethered nascent proteins are partially encapsulated upon GroES binding to GroEL, and refold in the chaperonin cavity. Reconstitution of chaperone competition at the ribosome shows that both Trigger factor and GroEL can be accommodated on long nascent chains, but GroEL and DnaK are mutually antagonistic. Our findings extend the role of GroEL/ES in *de novo* protein folding, and reveal an unexpected plasticity of the chaperonin mechanism that allows cotranslational substrate encapsulation.

## Introduction

The chaperonins are essential components of the cellular chaperone network in all domains of life (Balchin *et al*, 2016). They form large cage-like structures in which substrate proteins are temporarily encapsulated (Hayer-Hartl *et al*, 2016). The best characterised chaperonin is *E. coli* GroEL/ES. GroEL subunits assemble into two heptameric rings arranged back-to-back, which are transiently capped by heptameric GroES in an ATP-driven conformational cycle (Fig 1A) (Braig *et al*, 1994; Saibil *et al*, 2013). Apo GroEL binds folding intermediates on the inside of the open ring. ATP binding to GroEL then induces a conformational change that allows binding of GroES, which displaces the substrate into the newly-formed GroEL/ES cavity (Clare *et al*, 2012; Gruber & Horovitz, 2016; Hunt *et al*, 1996; Xu *et al*, 1997). Encapsulation isolates substrates from inter-molecular interactions, and in some cases modulates substrate folding (Hayer-Hartl *et al*, 2016). Obligate substrates often critically require the chaperonin to reach their native conformation on a physiologically-relevant timescale (Balchin *et al*, 2020; Georgescauld *et al*, 2014; Singh *et al*, 2020; Weaver *et al*, 2017).

**Figure 1.**
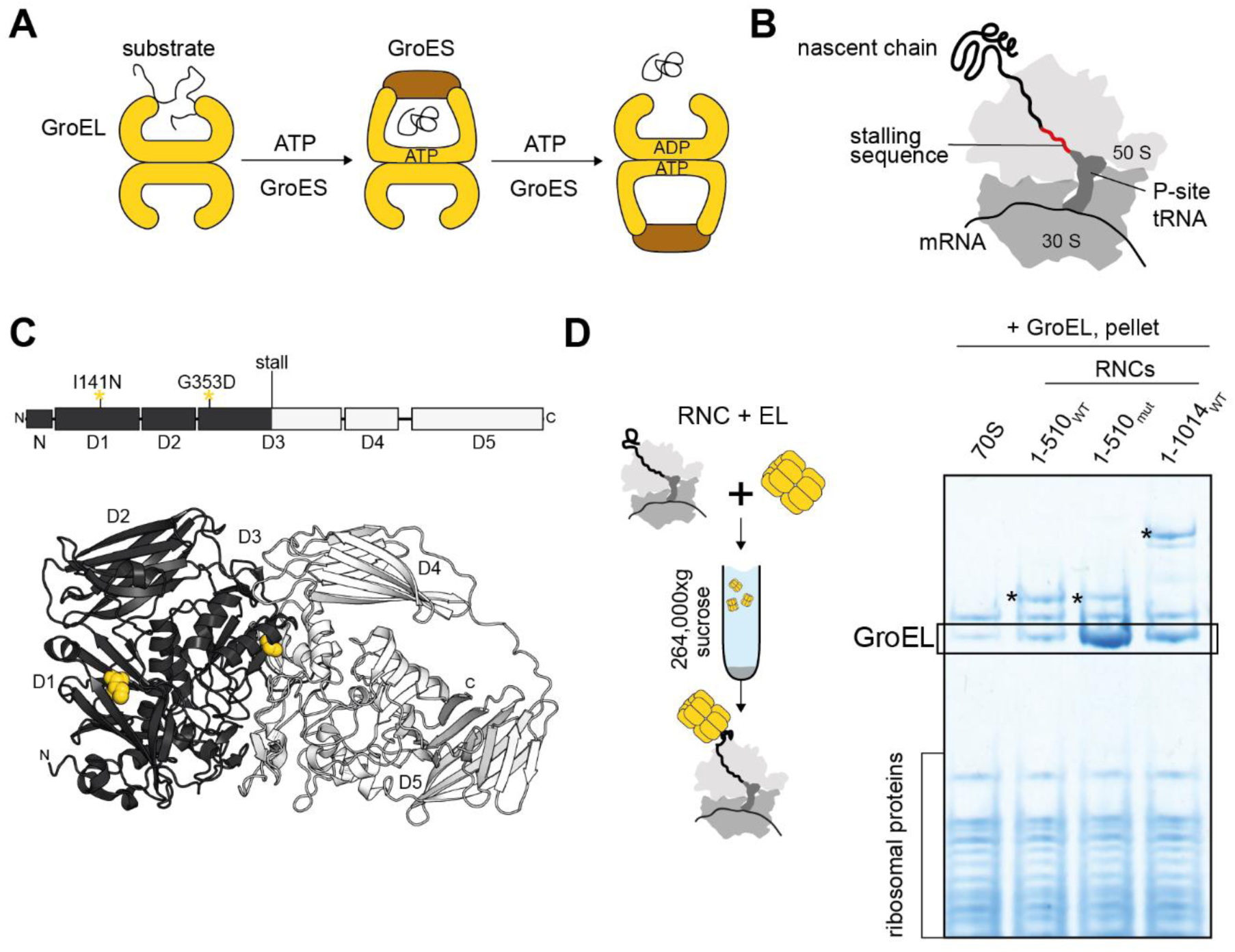
GroEL binds stalled ribosome:nascent chain complexes. **(A)** Simplified functional cycle of GroEL/ES. **(B)** Stalled 70S ribosome:nascent chain complex (RNC). **(C)** Domain organisation and monomer structure (PDB: 6CVM) of *E. coli* β-galactosidase (β-gal). Positions of destabilising mutations I141N and G353D are shown in yellow. The first 510 residues of β-gal, corresponding to RNC_1-510_, are colored black. **(D)** GroEL binds preferentially to a conformationally destabilized RNC. Left: schematic of co-sedimentation assay. Right: Coomassie-stained SDS-PAGE of the resuspended ribosomal pellet. Prior to sedimentation, GroEL was incubated with either empty ribosomes (70S), wild-type RNC_1-510_ (1-510_WT_), mutated (I141N, G353D) RNC_1-510mut_ (1-510_mut_), or RNC_1-1014_ (1-1014_WT_). RNCs were purified from Δ*tig* cells. Bands corresponding to the NCs (*) and GroEL are indicated.

GroEL/ES is part of a larger chaperone network that dynamically partitions between the ribosome and bulk cytosol. The most abundant components are Trigger factor (TF), which has direct affinity for ribosomes, and DnaK. GroEL overexpression can partially compensate for the loss of TF and DnaK in *E. coli* (Genevaux *et al*, 2004; Vorderwülbecke *et al*, 2004), suggesting partial functional redundancy. Although GroEL/ES most prominently functions post-translation and downstream of TF and DnaK (Langer *et al*, 1992), it also acts directly at the ribosome. GroEL binds translating ribosomes *in vitro* (Ying *et al*, 2006, 2005) and is recruited to a subset of nascent chains (NCs) *in vivo* (Zhao *et al*, 2021). At least 30 different NCs bind GroEL, representing ∼10% of all GroEL substrates (Fujiwara *et al*, 2010; Kerner *et al*, 2005; Zhao *et al*, 2021).

The post-translational function of GroEL/ES has been extensively characterized using chemically-denatured, full-length substrate proteins. Unlike these models, partially-synthesised NCs lack complete sequence information, are sterically influenced by proximity to the ribosome surface, and can populate unique structures compared to bulk solution (Agirrezabala *et al*, 2022; Chan *et al*, 2022; Kaiser *et al*, 2011; Roeselová *et al*, 2024; Wales *et al*, 2024). How GroEL binds and modulates cotranslational folding intermediates is unclear. Moreover, ribosome-tethered NCs substantially exceed the estimated ∼70 kDa size-limit of the chaperonin cavity (Xu *et al*, 1997), raising the question of whether cotranslational substrates can be encapsulated by GroEL/ES.

Here, we address the mechanism of chaperonin function during cotranslational folding. We show that ribosome-tethered polypeptides bind the inside of the GroEL cavity and are partially encapsulated by GroEL/ES. The NC is locally destabilised by GroEL binding, but recovers its original conformation upon encapsulation. Our data suggest a role for GroEL/ES in rescuing cotranslational misfolding, and indicate that substrates can benefit from confinement in the chaperonin cavity before they are released from the ribosome.

## Results

### GroEL binds stalled ribosome:nascent chain complexes

As cotranslational chaperone substrates, we prepared *E. coli* ribosome:nascent chain complexes (RNCs) stalled during synthesis of a well-characterised model protein, β-galactosidase (β-gal) (Fig 1B,C). We previously found that these RNCs co-purified with sub-stoichoimetric levels (∼0.5%) of GroEL (Roeselová *et al*, 2024), prompting us to reconstitute GroEL:RNC complexes using purified components *in vitro*. GroEL co-sedimented with several β-gal RNCs, indicative of stable complex formation (Fig 1D and S1). GroEL also co-sedimented with empty 70S ribosomes, albeit to a lesser extent, suggesting a weak interaction with ribosomes even in the absence of NC. GroEL binding to RNC_1-510_, exposing 2½ domains of β-gal, could be further stabilized by mutating the NC to interfere with native folding of domains 1 and 3 (Fig 1C,D). GroEL therefore interacts with both empty ribosomes and ribosome-associated NCs, and is sensitive to the conformation of the NC.

### GroEL uses different surfaces to bind ribosomes and nascent chains

GroEL is expected to be flexibly tethered to ribosomes via the partially-folded NC, resulting in a highly dynamic assembly. To characterise the topology of these assemblies, we crosslinked complexes between GroEL and 3 different RNCs using disuccinimidyl dibutyric urea (DSBU) and identified crosslink sites using mass spectrometry (Fig 2A). GroEL crosslinked extensively to the NC in all cases (Fig 2B and S2A,B). We also identified numerous crosslinks between GroEL and the ribosomal stalk, L7/L12, which coordinates elongation factors during translation (Liljas & Sanyal, 2018). Crosslinking to the stalk did not depend on NC binding, as the same crosslinks formed when GroEL was mixed with empty 70S ribosomes (Fig 2D). Furthermore, no crosslinks were detected between GroEL and ribosomes stripped of L7/L12 (Fig S2C,D). GroEL therefore engages both NCs and ribosomes, the latter potentially via the stalk complex.

**Figure 2.**
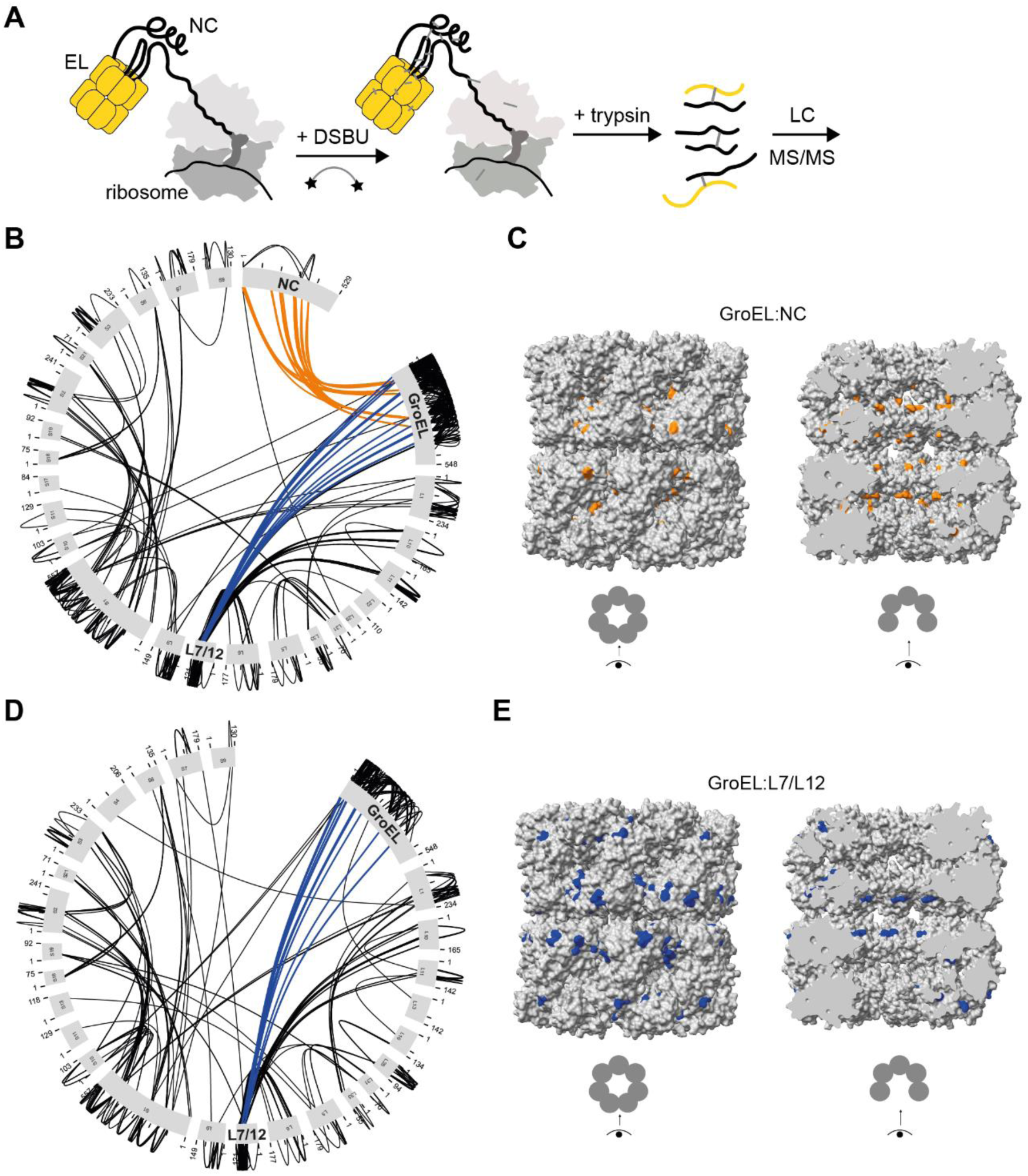
GroEL uses different surfaces to bind ribosomal proteins and nascent chains. **(A)** Crosslinking-mass spectrometry (XL-MS) experiment. **(B)** GroEL crosslinks extensively to both the NC and ribosomal stalk proteins. Map of crosslinks between GroEL and RNC_1-510mut_. Crosslinks between GroEL and the NC (orange) or L7/L12 (blue) are highlighted. **(C)** NCs crosslink to the inner surface of the GroEL cavity. Position of GroEL residues that crosslink to NC_1-510_, NC_1-510mut_, or NC_1-1014_ are shown in orange on the structure of GroEL ((Roh *et al*, 2017) PDB: 5W0S). Left, outer surface. Right, inner surface. **(D)** GroEL crosslinks to the L7/L12 stalk of empty ribosomes. Map of crosslinks between GroEL and empty 70S ribosomes. Crosslinks between GroEL L7/L12 (blue) are highlighted in blue. **(E)** L7/L12 crosslinks to the outer surface of GroEL. Position of GroEL residues that crosslink to L7/L12 in empty ribosomes, NC_1-510_, NC_1-510mut_, or NC_1-1014_ are shown in blue on the structure of GroEL (PDB: 5W0S). Left, outer surface. Right, inner surface.

Mapping the crosslink positions revealed a separation of binding interfaces. NCs preferentially crosslinked to residues inside the GroEL cavity, consistent with a substrate-like interaction (Fig 2C). In contrast, L7/L12 preferentially crosslinked to the outer surface of the GroEL (Fig 2E). GroEL co-sedimentation with RNC_1-510mut_ was unchanged after removing the L7/L12 stalk, showing that stalk binding is not strictly required for stable binding of GroEL to RNCs (Fig S2E).

### Nascent chains bind the apical domains and C-terminal tails of GroEL

We next focused on RNC_1-510mut_, which formed the most stable complex with GroEL among the tested RNCs (Fig 1D). To delineate the NC binding surface on GroEL, we measured RNC-induced protection of amide hydrogens using hydrogen/deuterium exchange-mass spectrometry (HDX-MS) (Fig 3A). Compared to isolated GroEL, GroEL bound to RNC_1-510mut_ was protected from deuterium exchange at 4 different sites per subunit (Fig 3B,C). The most protected site (residues 519-548) was the hydrophobic C-terminal tail of GroEL, which is structurally disordered and protrudes into the cavity (Fig 3C,D). A hydrophobic surface on the apical domain (residues 195-214) was also protected, near residues that crosslinked to NC_1-510mut_ (Fig 3C,E). This is consistent with previous studies showing that residues 199, 201, 203 and 204 are involved in substrate binding (Fenton *et al*, 1994), and cryo-EM structures which locate post-translational substrates at the apical domains of GroEL (Balchin *et al*, 2018; Clare *et al*, 2009; Elad *et al*, 2007; Gardner *et al*, 2023; Mamchur *et al*, 2021; Weaver *et al*, 2017). The two other protected sites (134-161, 454-488) are close together on the external surface of GroEL, near the ATP-binding site (Fig 3C,E). These peptides are close to residues that crosslink to L7/L12, suggesting that the protection in this region may be caused by interactions with the ribosome rather than NC.

**Figure 3.**
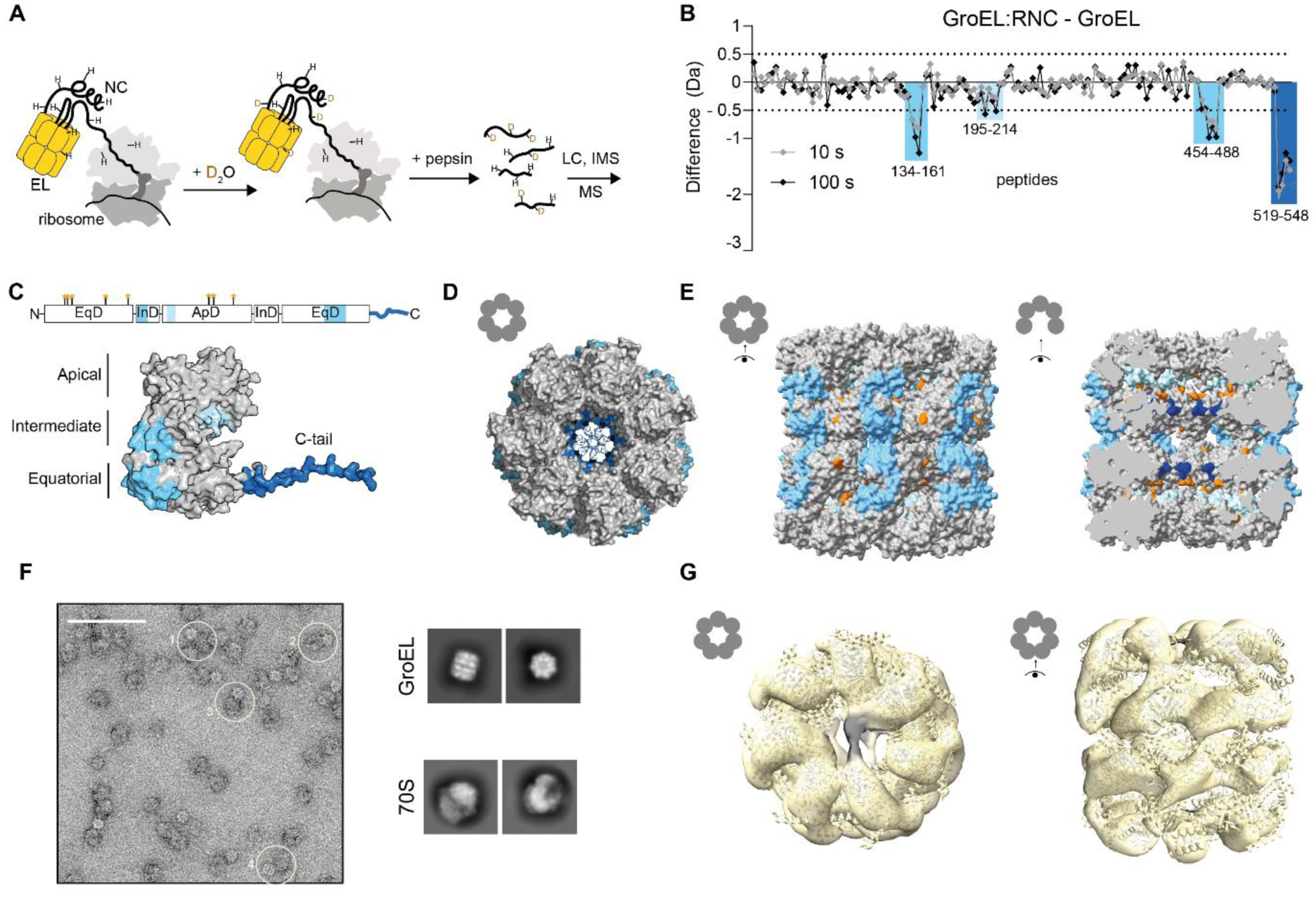
Nascent chains bind the apical domains and C-terminal tails of GroEL. **(A)** Hydrogen/deuterium exchange-mass spectrometry (HDX-MS) experiment. **(B)** Protection of GroEL induced by RNC binding. Difference in deuterium uptake after 10 s (grey) or 100 s (black), between GroEL bound to RNC_1-510mut_ and isolated GroEL. Values are plotted for individual GroEL peptides. Negative values indicate less deuteration of a peptide in RNC-bound GroEL relative to isolated GroEL, and sites with a difference in deuterium uptake > 0.5 Da are colored blue. **(C)** Sites of RNC-induced protection from deuterium exchange, mapped onto the domain organization and predicted structure of monomeric GroEL (AF-P0A6F5-F1). Crosslink sites for RNC1-510mut are indicated on the domain schematic with orange asterisks. **(D)** As in (C), shown on a top-view tetradecameric GroEL. The C-tails protrude into the central cavity. **(E)** As in (C), showing a side-view of tetradecameric GroEL (PDB: 5W0S). Left, outer surface. Right, inner surface. GroEL residues that crosslinked to NC_1-510mut_ are shown in orange. Sites with a difference in deuterium uptake > 0.5 Da are colored blue. **(F)** Visualization of GroEL:RNC assemblies. Left: Negative stain electron microscopy (nsEM) micrograph of DSBU-crosslinked GroEL:RNC_1-510mut_ assemblies. The scale bar corresponds to 100 nm. Examples of GroEL molecules positioned near ribosomes are circled (1-4). Right: 2D class averages of GroEL and 70S ribosomes. **(G)** NC density spans the GroEL cavity at the apical domains. 3D reconstruction of DSBU-crosslinked GroEL:RNC_1-510mut_ (map iv) from nsEM, viewed from the top and side. The structure of GroEL ((Roh *et al*, 2017) PDB: 5W0S) is docked into the map, and excess density corresponding to the NC is coloured black.

Negative stain electron microscopy (nsEM) of GroEL:RNC complexes showed GroEL positioned near ribosomes (Fig 3F and S3A). Single-particle analysis resulted in a low-resolution reconstruction of ribosome-associated GroEL, and fitting the solved structure of GroEL (Roh *et al*, 2017) into the map revealed additional density contacting the apical domains at the cavity opening (Fig 3G and S3B,C). This density was poorly resolved in an untreated sample (Fig S3B), but clearly spans the cavity in maps from DSBU-crosslinked samples (Fig 3G and S3B).

In summary, our XL-MS, HDX-MS and nsEM analyses indicate that the NC binds near the top of the GroEL cavity, as previously observed for chemically denatured substrate proteins. Moreover, we provide evidence that the disordered C-termini of GroEL, established to be important for substrate capture and folding but unresolved in structures (Dalton *et al*, 2015; Gardner *et al*, 2023; Ishino *et al*, 2015; Tang *et al*, 2008; Weaver & Rye, 2014), directly contact the NC.

### GroES can be accommodated on RNCs bound by GroEL

During chaperonin-mediated protein folding, ATP binding to GroEL induces a large conformational change that primes the chaperone for binding GroES, resulting in substrate encapsulation in the cis cavity (Fig 1A). As RNCs (> 2.5 MDa) substantially exceed the size limit of the GroEL/ES chamber (∼70 kDa, (Xu *et al*, 1997)), we sought to understand whether the same mechanism applies co-translation. We first tested the nucleotide dependence of GroEL binding to RNCs in the absence of GroES. Addition of either ADP, ATP, ADP/BeF_x_ or ATP/BeF_x_ reduced the amount of GroEL that co-sedimented with RNC_1-510mut_ (Fig S4A). The conformational changes triggered by nucleotide binding therefore weaken, but do not completely disrupt, the interaction between NCs and GroEL.

GroES can bind either ring of GroEL, resulting in a mixture of single-and double-capped complexes. To generate homogenous assemblies, we supplemented the reactions with ATP/BeF_x_, established to result in symmetric EL:ES_2_ complexes with both cavities capped by GroES (Taguchi *et al*, 2004) (Fig 4A). Pre-forming the symmetrically-closed EL:ES_2_ complex prevented either GroEL or ES from co-sedimenting with RNCs, demonstrating that NC binding requires access to the chaperonin cavity (Fig 4B S4A,B). In contrast, GroEL/ES co-sedimented with the RNC if GroEL and GroES were allowed to bind prior to addition of ATP/BeF_x_. The amount of co-sedimenting GroEL decreased only slightly upon addition of GroES and ATP/BeF_x_, from 9.8±0.9 to 6.7±1.5 GroEL subunits per ribosome (Fig 4C and Table S2).

**Figure 4.**
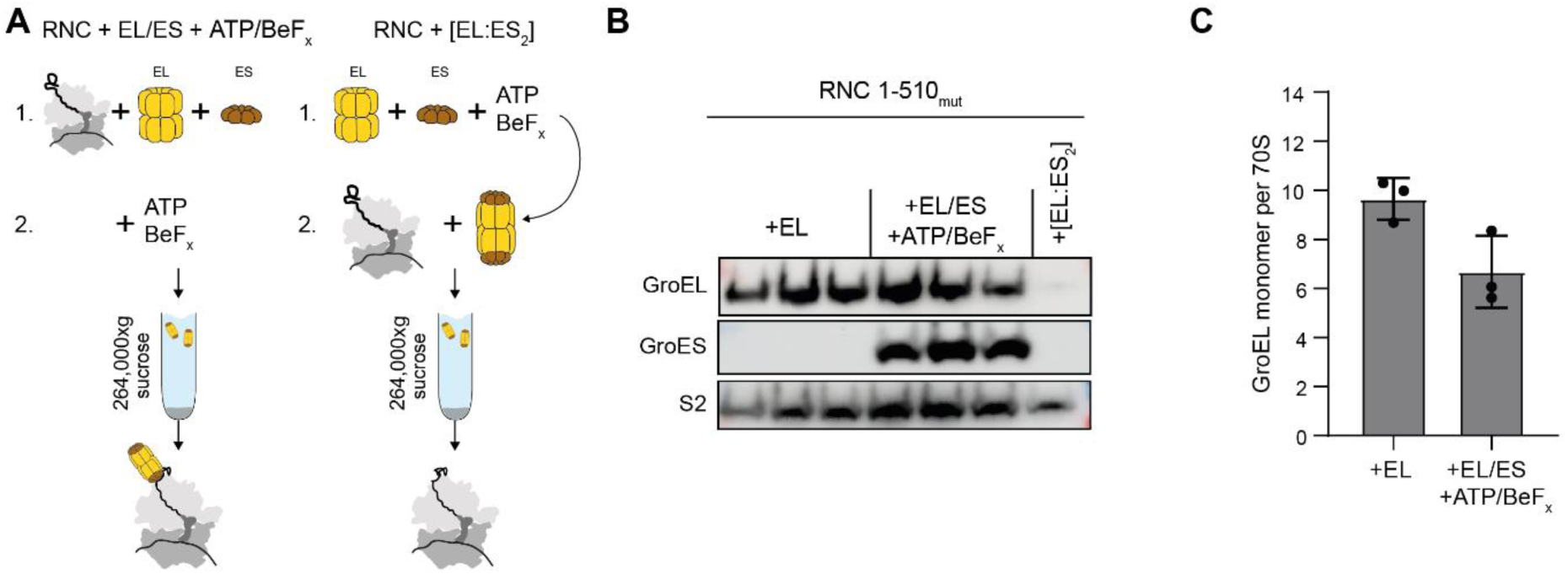
GroES can be accommodated on RNCs bound by GroEL. **(A)** GroEL/ES:RNC co-sedimentation assay. Two different orders of addition were followed. Left: RNCs were incubated with GroEL/ES then ATP/BeF_x_ was added. Right: GroEL/ES was pre-mixed with ATP/BeF_x_ before incubation with RNCs. **(B)** GroES co-sediments with a GroEL:RNC complex. Immunoblot of the resuspended ribosomal pellets from triplicate co-sedimentation assays described in (A), probed using antibodies against GroEL, GroES and the ribosomal protein S2. **(C)** Stoichiometry of GroEL:RNC complexes. Mean intensity-based absolute quantification (iBAQ) values for GroEL in resuspended ribosomal pellets from co-sedimentation assays described in (A). Values are normalised to the average iBAQ of all ribosomal proteins in each sample. Error bars represent the SD of three independent co-sedimentation assays.

### GroEL/ES partially encapsulates ribosome-bound nascent chains

As GroES could be stabilized on GroEL:RNC complexes, we next asked whether NCs are encapsulated by the chaperonin. We first used XL-MS to probe the architecture of the ATP/BeF_x_-stabilised EL:ES_2_:RNC complex. Similar to GroEL:RNC complexes, both GroEL and GroES crosslinked to the NC and ribosomal stalk (Fig 5A and S5A). Residues that crosslinked to the L7/L12 stalk mapped exclusively to the outside of the GroEL/ES, while residues that crosslinked to the NC were biased to the inside of the EL/ES cavity (Fig 5B and S5B). The latter included four residues (226, 360, 364, 371) that did not crosslink to NCs in GroEL:RNC complexes (Fig S2). Two of these residues (364, 371) had instead crosslinked to L7/L12 in the GroEL:RNC complexes (Table S3). The other two residues (226, 360) were previously shown to interact with substrate proteins in the closed EL:ES_2_ complex (Gardner *et al*, 2023). Mapping these residues onto the GroEL and EL:ES_2_ structures showed that they were positioned inside the GroEL cavity in the EL:ES_2_ complex, but not apo-GroEL, explaining the difference in crosslinking patterns (Fig S5C).

**Figure 5.**
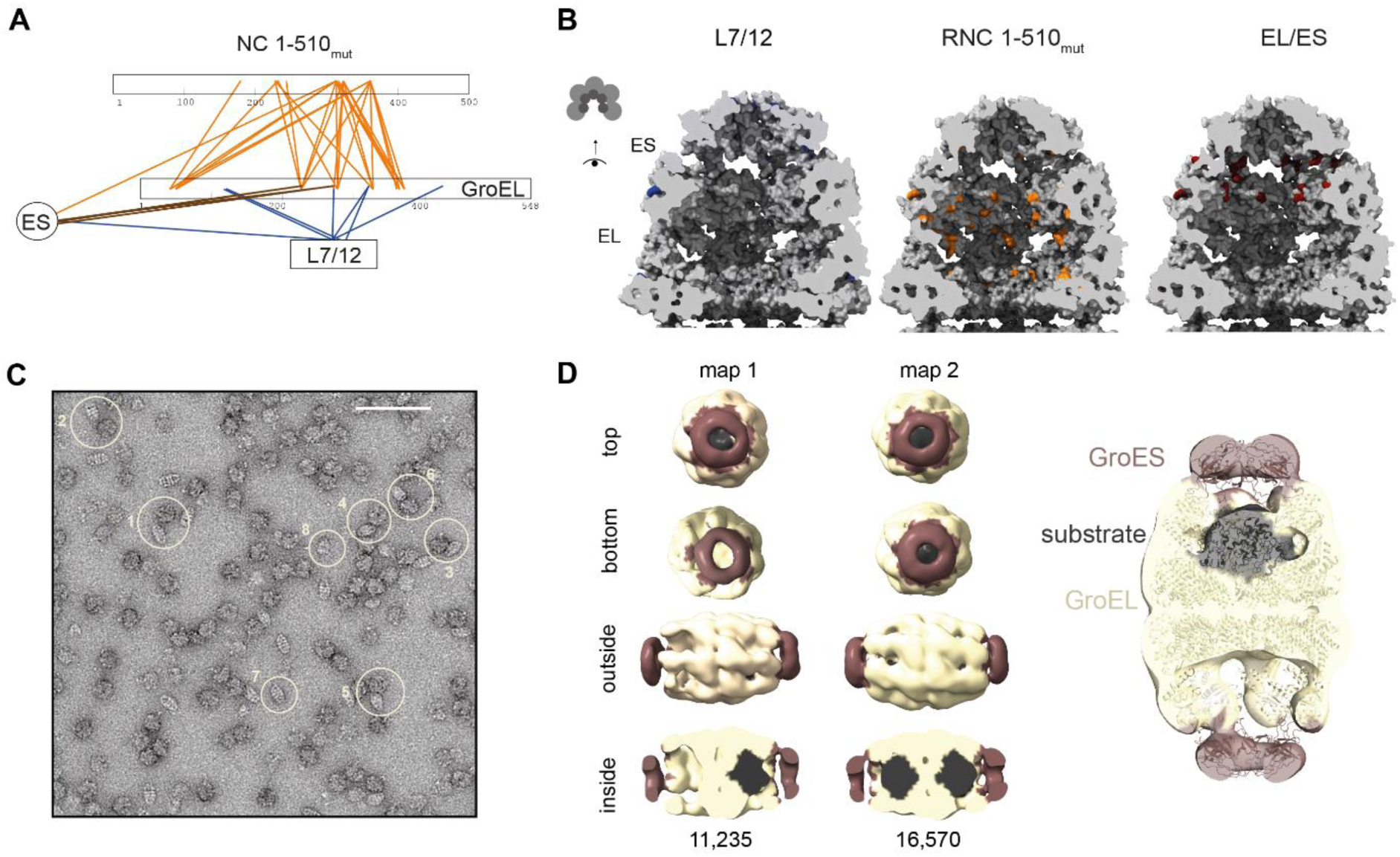
Nascent chains occupy the central cavity of GroEL/ES. **(A)** Map of crosslinks between GroEL/ES and RNC_1-510mut_ in the EL:ES_2_::RNC complex. Only crosslinks between GroEL/ES and L7/L12 (blue), GroEL and GroES (brown), and GroEL/ES and the NC (orange) are shown. **(B)** NC and ribosome stalk crosslink to the inner and outer surface of GroEL/ES, respectively. Crosslink sites are mapped onto the structures of the ATP/BeF_x_-stabilised GroEL/ES_2_ complex ((Kim *et al*, 2022) PDB:7VWX), cut away to show the inner surface of the GroEL/ES cavity. Residues are separated according to whether they crosslink to L7/L12 (blue), the NC (orange), or connect GroEL and GroES (brown). **(C)** Visualization of GroEL:ES_2_:RNC assemblies. Micrograph from nsEM of GroEL:ES_2_:RNC_1-510mut_ complex. The scale bar corresponds to 100 nm. Examples of GroEL:ES_2_ complexes positioned near ribosomes (1-6), isolated double-capped EL:ES_2_ (7), or single-capped EL:ES_1_ (8) complexes are circled. **(D)** NC density in the GroEL/ES cavity. Left: 3D reconstructions of GroEL:ES/RNC complexes from nsEM. Two classes can be distinguished, with additional density (black) in one or both chambers. The number of particles contributing to each reconstruction is given at the bottom. Right: The structure of GroEL/ES with encapsulated Rubisco ((Kim *et al*, 2022) PDB: 7VWX) docked into map 1.

The position of the nascent chain inside the GroEL/ES cavity was further supported by nsEM of the EL:ES_2_:RNC complex. GroEL/ES complexes were positioned near ribosomes (Fig 5C), and 90% were double-capped EL:ES_2_ (Fig S5D). Analysis of these species resulted in two 3D reconstructions which revealed additional density inside one or both EL/ES cavities (Fig 5D and Fig S5E).

As an orthogonal test of NC encapsulation, we treated RNCs with proteinase K (Fig 6 and S6A). Addition of either GroEL, or GroEL/ES followed by ATP/BeF_x_, to RNC_1-510mut_ slowed proteolysis of the full-length NC-tRNA species. Moreover, binding of EL:ES_2_ uniquely protected two ∼50-60 kDa proteolysis intermediates. These were truncated near the C-terminus of the NC, as they had lost the peptidyl tRNA and reacted with an antibody against the N-terminus of β-gal (Fig S6A). The same proteolysis intermediates were observed when EL:ES_2_:RNC complexes were isolated from excess unbound chaperone prior to protease treatment (Fig S6B,C). These data suggest that GroEL:ES_2_ can encapsulate a ∼60 kDa NC fragment prior to NC release from the ribosome, consistent with the estimated size limit of the GroEL/ES cavity.

**Figure 6.**
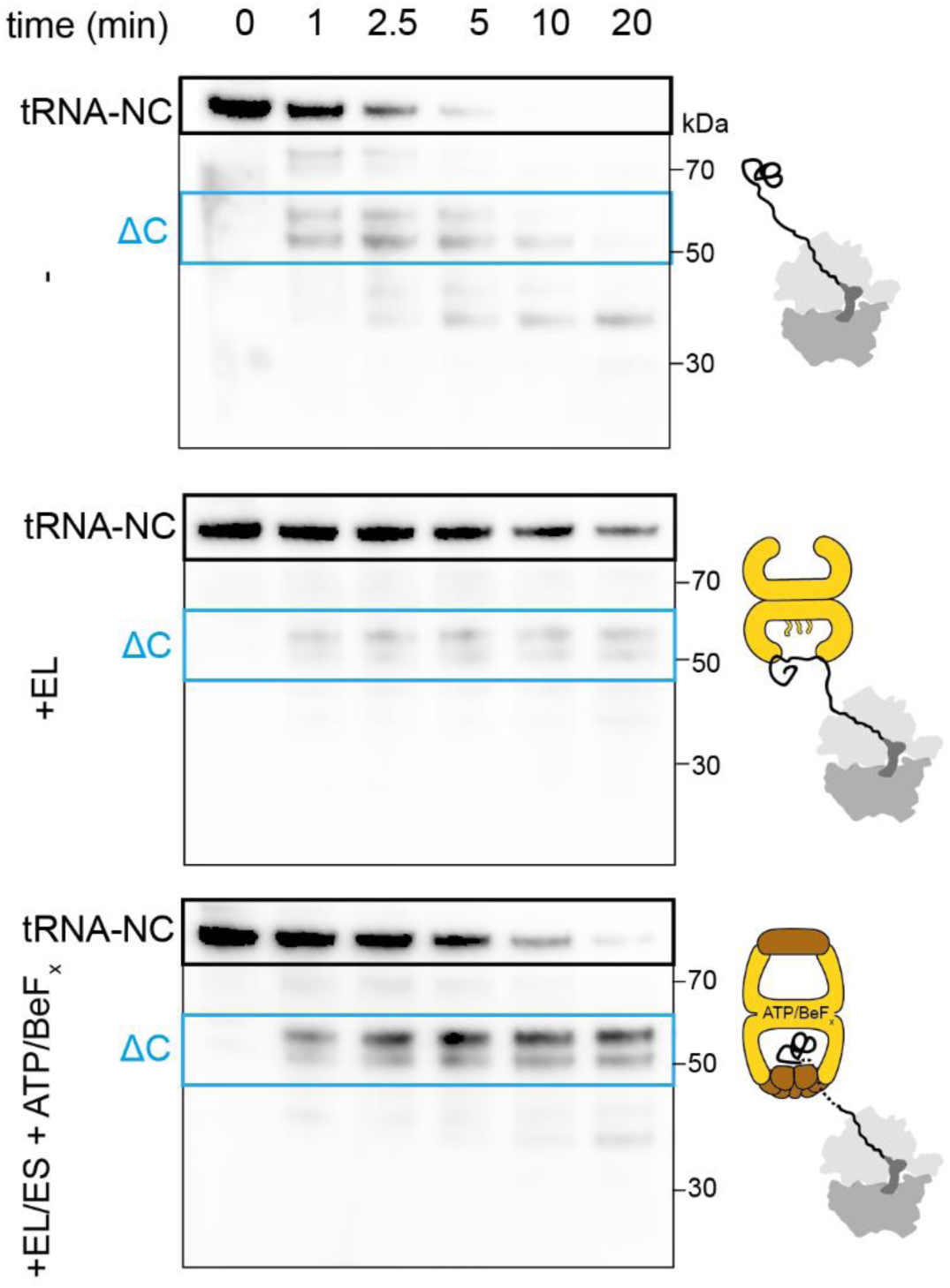
GroEL/ES protects ∼60 kDa fragments of nascent β-gal from limited proteolysis. NC_1-510mut_ at different times following addition of proteinase K, detected by immunoblotting using an antibody raised against full-length β-gal. Limited proteolysis was performed on isolated RNC_1-510mut_ (top), RNC_1-510mut_ incubated with GroEL (middle) and RNC_1-510mut_ incubated with GroEL and GroES followed by addition of ATP/BeF_x_ (bottom). Bands corresponding to the tRNA-bound full-length NC (tRNA-NC) and degradation intermediates truncated at the C-terminus (ΔC) are highlighted.

In summary, our XL-MS, nsEM and limited proteolysis data demonstrate that ribosome-tethered NCs are partially encapsulated by GroEL/ES.

### GroEL binding destabilises the NC prior to encapsulation

To understand how GroEL binding affects the conformation of the NC, we analysed GroEL:RNC and EL:ES_2_:RNC complexes using HDX-MS, and compared deuterium uptake to isolated RNC_1-510mut_. Peptide coverage of the nascent chain was limited to ∼55% by the extreme complexity of the sample containing both ribosome- and GroEL-derived peptides (Fig S7A,B). We found that the N-terminal part of the NC (residues 8-83) was deprotected by 0.5-1.5 Da upon GroEL binding, indicative of local conformational destabilisation (Fig 7A). Comparison of NC_1-510mut_ to native full-length β-gal showed that residues 8-24 were highly deprotected (> 4 Da) in the NC, while residues 25-83 showed near-identical deuterium uptake to the native state (Δ < 0.5 Da, Fig 7B). GroEL binding therefore destabilises a native-like region in the NC, and further destabilises an already partially unfolded segment.

**Figure 7.**
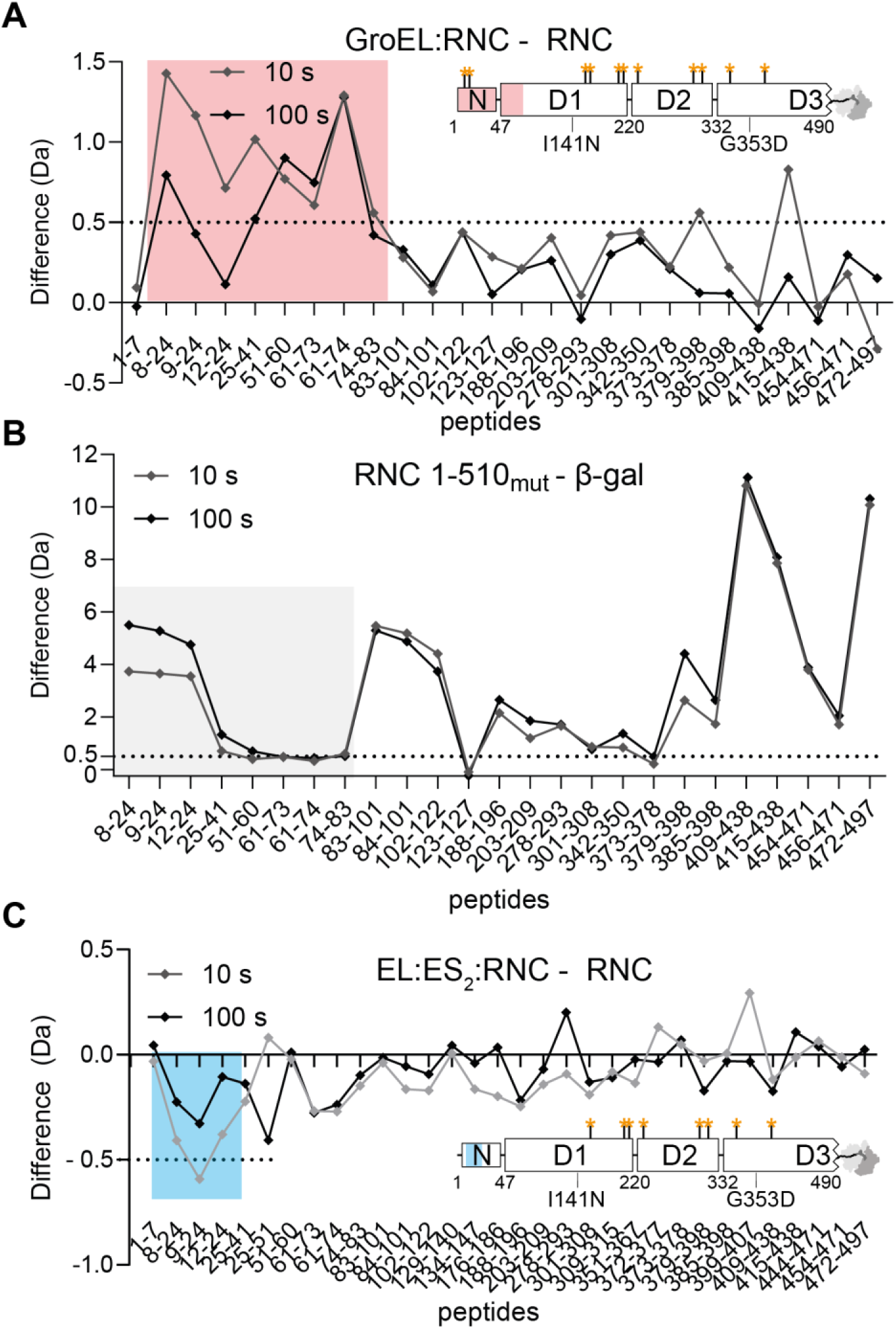
GroEL locally destabilizes the NC prior to encapsulation in GroEL:ES_2_. **(A)** Deprotection of NC_1-510mut_ upon binding GroEL. Difference in deuterium uptake after 10 s (grey) or 100 s (black), between GroEL-bound RNC_1-510mut_ and isolated RNC_1-510mut_. Values are plotted for individual peptides covering the β-gal NC. Positive values indicate more deuteration of a peptide in the GroEL-bound RNC relative to the isolated RNC. The data are also mapped onto a schematic domain diagram of NC_1-510mut_, with sites on the NC that crosslinked to GroEL indicated using orange asterisks. **(B)** Conformational dynamics of NC_1-510mut_ compared to native β-gal. Difference in deuterium uptake after 10 s (grey) or 100 s (black), between isolated RNC_1-510mut_ and native full-length β-gal. Positive values indicate more deuteration of a peptide in the RNC relative to native β-gal. **(C)** No deprotection of NC encapsulated by GroEL/ES. As in (A), except isolated RNC_1-510mut_ is compared to EL:ES_2_:RNC_1-510mut_. Negative values indicate decreased deuteration in the encapsulated compared to the isolated RNC.

Since GroEL selectively destabilised the N-terminal region of the NC, we next asked whether this region specifically binds the chaperone. We found that the N-terminus is not required for binding, as deletion of the first 24 or 81 residues did not disrupt the GroEL:RNC interaction (Fig S7C). Instead of uniquely binding the N-terminus, our data argue that GroEL binds multiple sites across the NC and thereby prevents the N-terminal region from making stabilising intramolecular contacts. This is consistent with our XL-MS analysis, which identified crosslinks to GroEL throughout the NC (Fig 7A).

GroEL-induced NC destabilisation was reversed upon encapsulation in EL:ES_2_ (Fig 7C). An N-terminal segment was slightly protected (∼0.5 Da) from deuterium uptake relative to isolated RNC_1-510mut_, and no other region differed in exchange, indicating that the NC recovered its original fold. The NC therefore interacts differently with GroEL compared to EL:ES_2_, consistent with the burial of hydrophobic sites on GroEL upon GroES binding (Hayer-Hartl *et al*, 2016). This difference was also apparent when we analysed deuterium uptake of GroEL itself (Fig S7D). The substrate binding sites in the apical domains of apo GroEL (195-214, Fig 3) were unaffected by NC binding to EL:ES_2_, consistent with burial of these sites at the interface with GroES (Xu *et al*, 1997). Furthermore, the flexible C-termini were only weakly protected (Δ∼0.4 Da) upon RNC binding to the EL:ES_2_, indicative of reduced involvement in substrate binding compared to apo GroEL (Fig 3B).

### GroEL competes with TF and DnaK

We previously showed that β-gal RNCs bind both DnaK and TF, with TF outcompeting DnaK at ribosome-proximal sites (Roeselova *et al*, 2024). We therefore asked whether GroEL binding to RNCs is influenced by TF and DnaK (Fig 8A). Since RNC_1-510_ is a poor TF substrate (Roeselová *et al*, 2024), we prepared RNC_1-180_, RNC_333-510_ and RNC_1-1014_ which bind both TF and GroEL when added individually (Fig 8B,C). For the shorter NCs, GroEL binding was completely outcompeted by equimolar TF (Fig 8B). In contrast, GroEL or GroEL/ES binding to RNC_1-1014_ was not affected by, nor did it affect, TF binding (Fig 8C). This suggests that TF and GroEL can occupy independent sites on NCs, with TF preferred at the ribosome-proximal site. Simultaneous addition of GroEL and DnaK led to reduced levels of both chaperones on RNC_1-510_ or RNC_1-510mut_ (Fig 8D). The same behaviour was observed in the presence of GroES and ATP/BeF_x_ (Fig S8A,B). GroEL and DnaK therefore act in a mutually antagonistic manner at the ribosome.

**Figure 8.**
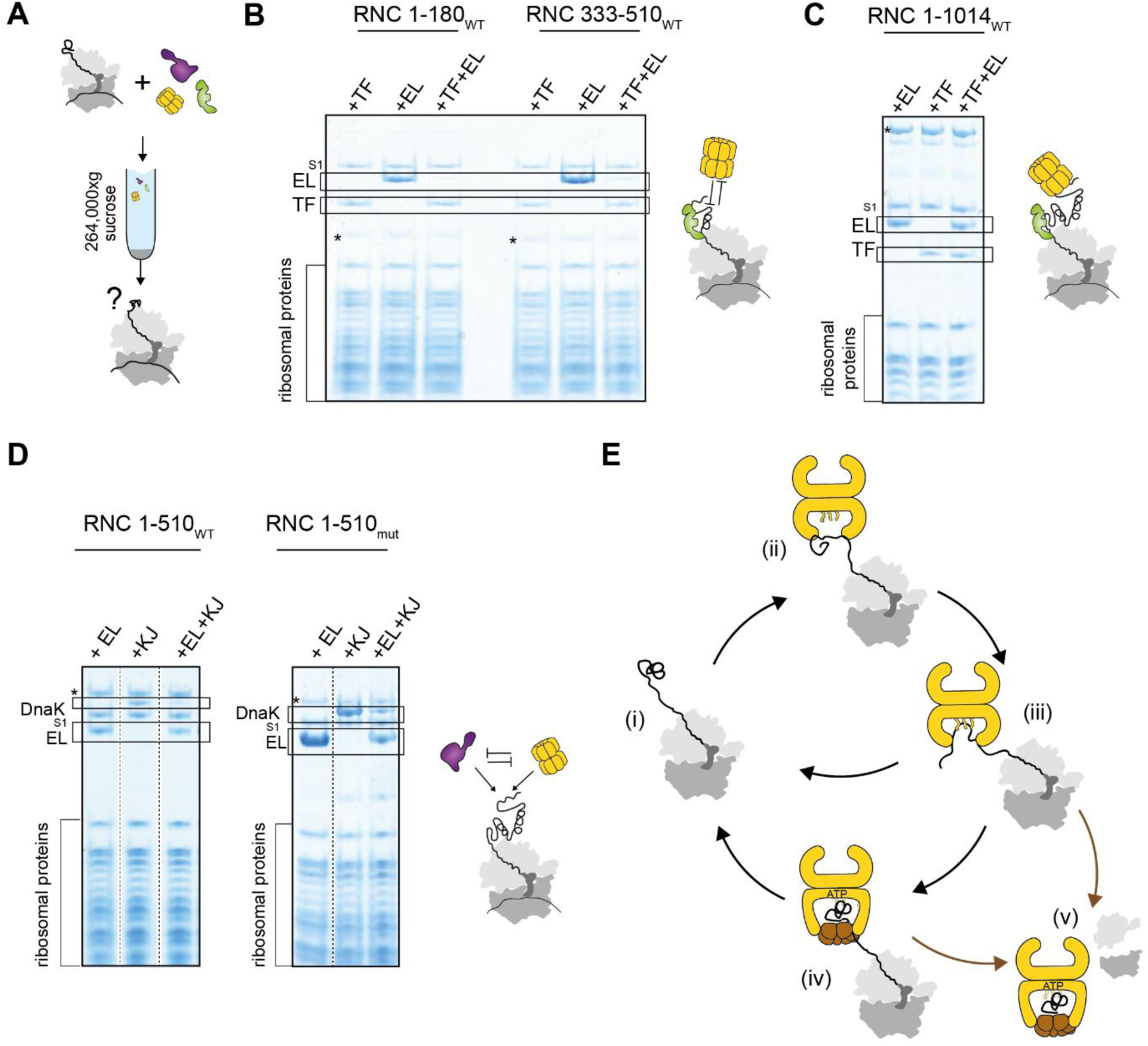
Coordination of GroEL with Trigger factor and DnaK. **(A)** Multi-chaperone co-sedimentation assay. RNCs were incubated with GroEL and either TF or DnaK/DnaJ/ATP before centrifugation through a 35% sucrose cushion to separate the ribosomal fraction (pellet) from unbound chaperones (supernatant). **(B)** TF outcompetes GroEL at short NCs. Coomassie-stained SDS-PAGE of the resuspended ribosomal pellet from the co-sedimentation assay. RNC_1-180_ or RNC_333-510_ were incubated with GroEL (+EL), Trigger factor (+TF), or both chaperones together (+EL+TF). Bands corresponding to the NC (*), TF and GroEL are indicated. **(C)** TF and GroEL do not compete for binding long NCs. Chaperone co-sedimentation with RNC_1-1014_ was tested as in (B). **(D)** GroEL and DnaK compete for binding RNCs. Coomassie-stained SDS-PAGE of the resuspended ribosomal pellet from the co-sedimentation assay as in (A). RNC_1-510WT_ or RNC_1-510mut_ were incubated either with GroEL (+EL), DnaK/DnaJ/ATP (+KJ) or both (+EL+KJ). Bands corresponding to the NC (*), DnaK and GroEL are indicated. **(E)** Model for the cotranslational action of GroEL/ES. Poorly folded NCs (i) are captured by the apical domains and C-terminal tails of GroEL (ii and iii), resulting in further unfolding. GroEL may dissociate, or be joined by GroES to partially encapsulate the ribosome-bound NC (iv). The NC may then fold enough to exclude GroEL/ES, or remain encapsulated and complete folding in the chaperonin cavity post-translation (v). Some substrates may only become encapsulated after translation termination (arrow from iii to v).

## Discussion

Our data provide a mechanistic underpinning for chaperonin function during cotranslational folding (Fig 8E). Despite the proximity of the ribosome, nascent polypeptides access binding sites in the GroEL cavity, including the apical domains and C-terminal tails. This interaction locally unfolds the NC, which then refolds following displacement into the cis cavity by GroES. GroEL recognises NCs that are poor TF substrates, and competes with DnaJ/K.

### Function of GroEL/ES at the ribosome

We show that GroEL/ES unfolds then partially encapsulates nascent polypeptides. GroEL-induced substrate expansion was previously observed, and is speculated to result from multivalent contacts with the chaperone cavity (Lin *et al*, 2008; Lin & Rye, 2004; Weaver *et al*, 2017; Weaver & Rye, 2014). In the context of cotranslational folding, local unfolding may resolve misfolded states prior to refolding in the GroEL/ES cage. Neither TF nor DnaK unfold β-gal NCs (Roeselova *et al*, 2024), suggesting that this may be a specialised function of GroEL.

The encapsulation of nascent chains is significant, as the chaperonin cavity is a unique folding environment which can modulate the energy landscape of protein folding. Physical confinement, the net negative charge of the cavity wall, and interactions with the disordered C-termini have all been shown to contribute to accelerated folding inside GroEL/ES (Hayer-Hartl *et al*, 2016). Furthermore, encapsulated substrates are insulated from intermolecular interactions, allowing folding at effectively infinite dilution. Our data show that nascent proteins can benefit from folding within the chaperonin cage before their synthesis is complete.

### Mechanism of partial encapsulation of nascent polypeptides

Cotranslational encapsulation implies that the NC protrudes from the closed chaperonin cavity to remain connected to the ribosome. This represents a topological problem, since GroES is thought to bind *en bloc* to GroEL and seal the cavity. One possibility is that GroES binds asymmetrically to GroEL, leaving a gap through which the NC can thread. Indeed, previous work has suggested that substrates might escape through a space at the GroEL/ES interface (Motojima & Yoshida, 2010). Furthermore, a recent structure of GroEL bound to Rubisco and ADP/BeF_3_ showed a subset of GroEL-domains in a GroES-binding state, suggesting that partial docking of GroES on GroEL is possible (Gardner *et al*, 2023). Understanding exactly how encapsulation is achieved will require structural resolution of the nascent polypeptide, a significant challenge due to the conformational heterogeneity of partially-folded substrates.

This mechanism may also be relevant post-translation. A fraction of obligate GroEL/ES substrates are too large (>70 kDa) to be completely encapsulated (Chaudhuri *et al*, 2009, 2001; Kerner *et al*, 2005; Paul *et al*, 2007; Sakikawa *et al*, 1999), and many large proteins are efficiently refolded by GroEL/ES (To *et al*, 2022). This has been proposed to occur without cis encapsulation, via iterative binding and release from the trans ring of GroEL (Chaudhuri *et al*, 2001). Our data suggest the alternative possibility that some large GroEL substrates fold via domain-wise encapsulation in the cis cavity. Indeed, a segmental encapsulation mechanism has previously been demonstrated for the eukaryotic chaperonin TRiC (Rüßmann *et al*, 2012). Whether TRiC can also encapsulate ribosome-tethered domains is not clear.

### GroEL binding to the ribosome

We show that GroEL weakly interacts with ribosomes independent of the NC. This occurs via the outer surface of the chaperonin cavity, via sites distinct from those that bind substrates. Although XL-MS suggests that GroEL binds directly to the ribosomal L7/L12 stalk, we do not exclude that a different interface is involved. For example, GroEL binding to ribosomal RNA would not be detected by DSBU crosslinking.

Whether GroEL binding to the ribosome influences the cotranslational activity of the chaperonin remains to be determined. Weak interactions with the ribosome may promote NC capture by increasing the local concentration of GroEL, or contribute avidity for NCs that are poor GroEL substrates. Ribosome binding may also increase the “processivity” of GroEL/ES, by keeping GroEL close to the ribosome between ATP-driven cycles of NC binding and release.

The L7/L12 stalk is involved in recruiting elongation factors to the ribosome (Diaconu *et al*, 2005; Liljas & Sanyal, 2018). If GroEL does bind directly to this site, it raises the intriguing possibility that GroEL may slow translation elongation during chaperonin-assisted cotranslational folding.

### Chaperone competition at the ribosome

Although GroEL may in principle recognise many different nascent polypeptides, competition with other cytosolic chaperones restricts GroEL to a subset of NCs *in vivo*. Indeed, simultaneous deletion of DnaK/DnaJ and TF was found to increase the level of GroEL at ribosomes ∼10-fold (Zhao *et al*, 2021). We show that TF efficiently outcompetes GroEL for binding to shorter NCs, but both chaperones can be accommodated on longer NCs. Longer NCs are also substrates of DnaK/J (Roeselová *et al*, 2024), resulting in competition between GroEL and DnaK. Considering that GroEL strongly prefers conformationally destabilised NCs, it may be recruited to nascent domains that fail to fold while they are close to the ribosome surface, and thus persistently expose hydrophobic segments beyond the reach of TF.

## Methods

### DNA vectors and cloning

The pET28-based plasmid encoding full-length β-galactosidase and the pET21-based plasmids encoding muGFP-tagged (Scott *et al*, 2018) RNC complexes stalled with SecM-based WWWPRIRGPP stalling sequence (Cymer *et al*, 2015) were cloned in our previous study (Roeselova *et al*, 2024). *E. coli* DnaK and DnaJ were expressed without any tags from a pET11d-based vector. His_6x_-tagged TF was expressed from a ProEX backbone. GroEL and GroES were expressed from pET-17b-based vectors (Novagen). Additional deletions and point mutations were introduced using site-directed mutagenesis with Q5 or Phusion polymerases (NEB). All constructs used in this study (Table S4) were verified by sequencing.

### RNC buffers

RNC low-salt buffer contained 50 mM HEPES-NaOH pH 7.5, 12 mM Mg(OAc)_2_, 100 mM KOAc, 1 mM DTT and 8 U/mL RiboLock RNase inhibitor (ThermoScientific). RNC high-salt sucrose cushion contained 35% sucrose, 50 mM HEPES-NaOH pH 7.5, 12 mM Mg(OAc)_2_, 1 M KOAc, 1 mM DTT, 8 U/mL RiboLock RNase inhibitor and 0.2x Halt Protease Inhibitor Cocktail (ThermoScientific). RNC low-salt sucrose cushion contained 35% sucrose, 50 mM HEPES-NaOH pH 7.5, 12 mM Mg(OAc)_2_, 100 mM KOAc, 1 mM DTT, 8 U/mL RiboLock RNase inhibitor and 0.2x Halt Protease Inhibitor Cocktail.

### Protein purification

RNCs, full-length β-galactosidase, DnaK, DnaJ and Trigger factor were expressed and purified as described previously (Roeselova *et al*, 2024). Sequences of purified proteins are listed in Table S4. RNC_1-510mut_ contained two destabilising mutations – previously characterised I141N (Plata *et al*, 2010) and G353D designed using DynaMut2 (Rodrigues *et al*, 2021).

GroEL (V381W, A384W, V387W) was overexpressed in *E. coli* Δ*lac* cells (Didovyk *et al*, 2017) and induced at 37°C with 1 mM IPTG overnight. Cells were harvested (4,000 g, 20 min) and resuspended in Buffer L (30 mM Tris-HCl pH 7.5, 30 mM NaCl, 1 mM EDTA, 1 mM DTT) with lysozyme, universal nuclease (Thermo Fisher), and cOmplete EDTA-free protease inhibitor cocktail. Following lysis by sonication, the soluble fraction was isolated through centrifugation (50,000 g, 1 h, 4°C) and loaded onto a self-packed DEAE column equilibrated in Buffer L. Peak fractions eluted with an NaCl gradient in Buffer L were combined, buffer exchanged into Buffer L, and loaded onto a heparin column equilibrated in Buffer L. Peak fractions eluted with an NaCl gradient were combined, diluted 3-fold with Buffer L, and loaded onto Resource Q column preequilibrated in Buffer L. Peak fractions eluted with an NaCl gradient were combined, diluted 3-fold with Buffer L, and loaded onto a heparin column preequilibrated in Buffer L. Peak fractions eluted with an NaCl gradient were combined, concentrated, and purified further using a Superose 6 size-exclusion column. Protein concentration was determined by absorbance using an extinction coefficient of 21,430 M^-1^ cm^-1^ and a molecular weight of 57.6 kDa.

GroES (L49W) with an N-terminal double StrepII-tag was overexpressed in *E. coli* BL21(DE3) cells (NEB) and induced at 37°C with 1 mM IPTG overnight. Cells were harvested (4,000 g, 20 min) and resuspended in Buffer L with lysozyme, universal nuclease, and cOmplete EDTA-fee protease inhibitor cocktail. Following lysis by sonication, the soluble fraction was isolated through centrifugation (50,000 g, 1 h, 4°C) and loaded onto a StrepTactin-II column. Peak fractions eluted with 2 mM d-desthiobiotin in Buffer L. The affinity tag was cleaved by overnight incubation with GST-3C protease. The sample was concentrated and purified further on a Superdex 200 column. Excessive 3C protease was removed using a GSTrap column. Protein concentration was determined by absorbance using an extinction coefficient of 6,690 M^-1^ cm^-1^ and a molecular weight of 10.5 kDa.

### Depletion of L7/L12 from ribosomes

*E. coli* 70S ribosomes or RNCs were depleted of the ribosomal stalk L7/L12 proteins using an adapted protocol based on NH_4_Cl/ethanol treatment (Savelsbergh *et al*, 2005). Complete 70S ribosomes (NEB) or RNCs (33 µL, 13.3 µM) were mixed and incubated (4 °C, 10 min) with 450 µL of ice-cold stripping buffer (20 mM Tris-HCl, pH 7.5, 0.6 M NH_4_Cl, 20 mM MgCl_2_, 5 mM β-mercaptoethanol, RiboLock RNase inhibitor). Subsequently, 250 µL of ice-cold ethanol was added to the sample, mixed and incubated (4 °C, 10 min) followed by a second addition of 250 µL of ice-cold ethanol and incubation (4 °C, 5 min). Sample was then layered over a low-salt sucrose cushion and centrifuged (4 °C, 264,000xg, 2 h) to isolate the ribosomal fraction. The pellet containing 70S ribosomes or RNCs depleted of the ribosomal stalk L7/L12 proteins was resuspended in RNC low-salt buffer and the depletion was confirmed by immunoblotting. For RNCs, integrity of the NC linked to peptidyl-tRNA was confirmed by SDS-PAGE and Coomassie staining.

### Proteinase K assay

RNC_1-510mut_ was diluted to 0.25-3 µM in RNC low-salt buffer and incubated (30 min, 30 °C) either alone, with 5-fold molar excess GroEL 14-mer, or with 5-fold molar excess GroEL 14-mer and 10-20-fold molar excess GroES 7-mer. The sample with GroES was subsequently supplemented with ATP/BeF_x_ (1 mM ATP, 1 mM BeSO_4_ and 10 mM NaF) and incubated for further 10 min at 30 °C. Samples were then cooled down to 4 °C. Optionally, to isolate only the ribosomal fraction, samples were separated in a sucrose cushion centrifugation and the ribosomal pellets were resuspended in RNC low-salt buffer supplemented with ATP/BeF_x_. Subsequently, a reference aliquot (t_0_) was removed from the samples (total or only the isolated ribosomal pellets) before initiating degradation of RNCs (0.2 µM) with Proteinase K (Millipore, 2.5 ng/μL) at 4 °C. Aliquots of reactions were removed and quenched at different times by 1:1 mixing with 5 mM PMSF in RNC low-salt buffer, and analysed by SDS-PAGE and immunoblotting. Anti-β-galactosidase antibody (abcam, ab221199) was used to detect all β-galactosidase fragments. Anti-β-galactosidase antibody (abcam, ab106567) was used to specifically detect β-galactosidase fragments with intact N-terminus (i.e. with C-terminal truncations).

### Co-sedimentation assays

For co-sedimentation assays, 1 μM RNCs or empty 70S ribosomes (NEB) were incubated (30 min, 30 °C) with (co-)chaperones (DnaK – 2-5 µM, TF – 2-5 µM, DnaJ – 1-2 µM, GroEL 14-mer – 2-5 μM, GroES 7-mer – 10-20 μM) in RNC low-salt buffer. For DnaK co-sedimentation assays the buffer was supplemented with 1 mM ATP. For GroEL/ES co-sedimentation assays, RNCs were first incubated with GroEL and/or GroES (20 min, 30 °C) followed by, when indicated, addition of 1 mM ADP or ATP optionally with 1 mM BeSO_4_ and 10 mM NaF (10 min, 30 °C) as described in figure captions. Pre-formed [EL/ES_2_] complex was prepared by incubating (10 min, 30 °C) GroEL and GroES in RNC low-salt buffer with at least 1 mM ATP, 1 mM BeSO_4_ and 10 mM NaF. Following incubation of ribosomes or RNCs with (co-)chaperones, the samples were loaded onto a 35% sucrose cushion, and pelleted by centrifugation (264,000 g, 2 h). Following two washes with ice-cold RNC low-salt buffer, the pellet was resuspended at 4 °C. Resuspended pellets from co-sedimentations assays were analysed by SDS-PAGE or proteomics analysis and where indicated used as input for other assays. The sucrose cushion and washing buffer were supplemented with ADP, ATP, BeSO_4_ or NaF where appropriate.

### Immunoblotting

Following SDS-PAGE, proteins were transferred onto a PVDF membrane using a Trans-Blot Turbo Transfer System (BioRad). Membranes were blocked in PBS-Tween with 5% non-fat milk for 1 hour at RT and incubated with appropriate primary antibodies (1:1,000 dilution in PBS-Tween with 5% non-fat milk) for 1 hour at RT. Following three washes (5 min, RT) with PBS-Tween, membranes were incubated with appropriate HRP-conjugated secondary antibodies (1:10,000 dilution in PBS-Tween with 5% non-fat milk) for 1 hour at RT and washed (3×5 min, RT). Membranes were developed by enhanced chemiluminescence using SuperSignal West Pico PLUS Chemiluminescent Substrate (ThermoScientific). Antibodies used for each immunoblot are specified in figure legends. Primary antibodies used in this study are mouse anti-GroEL (abcam, ab82592), rabbit anti-GroES (Enzo, ADI-SPA-210-D), rabbit anti-S2 (Antibodies-Online, ABIN2938988), rabbit anti-L7/L12 (abcam, ab225681), rabbit anti-β-galactosidase raised against the full-length protein (abcam, ab221199) and chicken anti-β-galactosidase raised against a 17 amino acid peptide from near the N-terminus (abcam, ab106567). Primary antibodies were detected with HRP-conjugated goat anti-rabbit (abcam, ab205718), anti-mouse (abcam, ab205719), and anti-chicken (abcam, ab97135) secondary antibodies.

### Proteomic quantification of protein content in ribosomal pellets

Protein content in resuspended pellets following co-sedimentation assays was determined as previously described (Roeselova *et al*, 2024) with slight modifications. In short, 10 µg of total protein estimated from the ribosome concentration (based on absorbance at 260 nm) was separated in 8 mm on NuPAGE Bis-Tris gels (ThermoScientific, 1.0 mm, 10-12 wells, 12%) followed by Quick Coomassie Stain (Neo Biotech) staining, band excision and destaining in extraction buffer (50% acetonitrile, 100 mM ammonium bicarbonate, 5 mM DTT, 16 h, 4 °C). Samples were subsequently alkylated (40 mM chloroacetamide, 160 mM ammonium bicarbonate, 10 mM TCEP, 20 min, 70 °C), dehydrated in 100% acetonitrile, air-dried, and digested with trypsin (Promega). Tryptic peptides were loaded onto Evotips (Evosep) and eluted using the 30SPD gradient via an Evosep One HPLC (Bache *et al*, 2018) with a 15 cm C18 column into a Lumos Tribrid Orbitrap mass spectrometer (ThermoScientific) via a nanospray emitter (2,200 V). Acquisition parameters were set to data-dependent mode with precursor ion spectra acquired at 120,000 resolution followed by higher energy collision dissociation. Raw files were processed in MaxQuant (Cox & Mann, 2008) and Perseus (Tyanova *et al*, 2016) with Uniprot *E. coli* reference proteome database and a database for common contaminants. Protein and peptide false detection rates using a decoy reverse database were set to 1%. Quantification of proteins was achieved using iBAQ (intensity-based absolute quantification) and values were normalised to the average intensity of 70S ribosomal proteins. Statistical analyses was performed in GraphPad Prism 9. Raw and normalised values are listed in Table S2.

### Equilibrium HDX-MS analysis of RNCs

HDX-MS analysis of RNC:chaperonin complexes was conducted as previously described (Roeselova *et al*, 2024) with slight modifications. In short, stocks of freshly purified RNCs and RNC:chaperonin complexes (prepared via cosedimentation assays) were diluted to 5-6 µM in RNC low-salt buffer. Additionally, stocks of full-length native β-galactosidase, empty 70S ribosomes (NEB), GroEL and pre-closed ATP/BeF_x_-stabilised [EL:ES_2_] complex were prepared as controls. Deuterium labelling was initiated by 1:10 dilution of the stock solution in deuteration buffer (10 mM HEPES-NaOD, pD 7.5, 30 mM KOAc, 12 mM Mg(OAc)_2_, 1 mM DTT, RiboLock RNase inhibitor, 97% D_2_O). Following labelling at 25 °C for 10 or 100 seconds, the reaction was quenched with an equal volume of ice-cold quench buffer (100 mM sodium phosphate, pH 1.4, 4 M guanidium hydrochloride, 10 mM TCEP) lowering the pH to 2.5. Digestion was performed using agarose-immobilised pepsin (100 s, 10 °C). The sample was then filtered (0.22 µm PVDF filters, 13,000 g, 15 s, 0 °C) and snap frozen in liquid nitrogen for short-term storage. The same protocol was followed to prepare undeuterated controls, except the deuteration buffer was replaced by a H-based buffer (10 mM HEPES-NaOH in H_2_O, pH 7.5, 30 mM KOAc, 12 mM Mg(OAc)_2_, 1 mM DTT, RiboLock RNase inhibitor). Note that the RNC low-salt buffer and deuteration buffer were supplemented with 1 mM ATP, 1 mM BeSO_4_ and 10 mM NaF when labelling the EL:ES_2_:RNC complex and relevant control samples.

Frozen samples were thawed and injected into an Acquity UPLC M-class system with the cooling chamber containing the chromatographic columns kept at 0 ± 0.2 °C. Peptides were trapped (4 minutes, 200 µL/min) on a C4 trap column (Acquity BEH C4 Van-guard pre-column, 2.1 mm X 5 mm, 1.7 μm, Waters) and separated on a reverse phase Acquity UPLC HSS T3 column (1.8 µm, 1 mm X 50 mm, Waters) at a flow rate of 90 µL/min using a 25 min 3-30% gradient of acetonitrile in 0.1% formic acid. Analysis was performed using a Waters Synapt G2Si HDMS^E^ instrument in ion mobility mode, acquiring in positive ion mode over a range of 50 to 2,000 m/z with the conventional electrospray ionisation source operated at a source temperature of 80 °C with the capillary set to 3 kV.

MS^E^ data were processed using Protein Lynx Global Server (PLGS, Waters) to identify peptides in the undeuterated control samples using information from a non-specific cleavage of a database containing sequences of *E. coli* β-galactosidase, GroEL, GroES, 70S ribosomal proteins as well as porcine pepsin. PLGS search was performed using energy thresholds of low = 135 counts and elevated = 30 counts. Peptides identified by PLGS were subsequently filtered and processed in DynamX (Waters) with filters of minimum products per amino acid of 0.05 and minimum consecutive products of 1. All spectra were manually inspected, and poor-quality assignments were removed. Relative deuterium uptake in Da was calculated by subtracting the centroid mass of undeuterated peptides from those of deuterated peptides. Fractional uptake was calculated by dividing the relative uptake by the theoretical maximum for each peptide, equal to n-1, where n is the peptide length excluding prolines. Mean values of deuterium uptake are reported as relative as they are not corrected for back-exchange. Uptake differences >0.5 Da at any time point were considered to be meaningful. All uptake data, peptide coverage maps, and a summary of experimental conditions (Masson *et al*, 2019) are shown in Table S1.

### Crosslinking mass spectrometry

Crosslinking of ribosome:chaperonin or RNC:chaperonin complexes was conducted as described previously (Roeselova *et al*, 2024) with slight modifications. Purified GroEL and GroES were first buffer-exchanged into RNC low-salt buffer using Micro Bio-Spin 6 Columns (Biorad) to remove any traces of Tris-HCl. Empty ribosomes (NEB) or RNCs freshly purified from *Δtig E. coli* were subsequently diluted in RNC low-salt buffer to a final concentration of 1-2 µM and incubated (30 min, 30 °C) with 5 µM GroEL with or without 10 µM GroES. Samples with GroES were additionally incubated with 1 mM ATP, 1 mM BeSO_4_ and 10 mM NaF (10 min, 30 °C). Complexes were then cooled to 25 °C and 1 mM DSBU (ThermoScientific) was added to initiate the crosslinking reaction (1 h, 25 °C) followed by quenching with 20 mM Tris-HCl, pH 7.5. Finally, the ribosomal fraction was pelleted via sucrose cushion centrifugation (264,000 g, 2 h, 4 °C) and resuspended in RNC low-salt buffer.

Crosslinked samples were subsequently reduced (10 mM DTT), alkylated (50 mM iodoacetamide) and digested with trypsin. Tryptic peptides were fractionated using a high pH reverse phase chromatography (gradient of acetonitrile in 10 mM NH_4_HCO_3_, pH 8, TARGA C18 columns, Nest Group Inc.), lyophilized, and resuspended in 1% formic acid and 2% acetonitrile. Samples were then subjected to nano-scale capillary LC-MS/MS using the Vanquish Neo UPLC (ThermoScientific Dionex), a C18 PepMap Neo nanoViper trapping column (5 μm, 300 μm × 5 mm, ThermoScientific Dionex), and an EASY-Spray column (50 cm × 75 μm ID, PepMap C18, 2 μm particles, 100 Å pore size, ThermoScientific). Peptides were eluted with a gradient of acetonitrile over 90 minutes. Analysis was performed using a quadrupole Orbitrap mass spectrometer (Orbitrap Exploris 480, ThermoScientific) with a nano-flow electrospray ionisation source. Acquisition parameters were set to data-dependent mode using a top 10 method, recording a high-resolution full scan (R=60,000, m/z 380-1,800) followed by higher energy collision dissociation (stepped collision energy 30 and 32% Normalized Collision Energy) of the 10 most intense MS peaks, excluding ions with precursor charge state of 1+ and 2+. The fragment ion spectra were acquired at a resolution of 30,000 and a dynamic exclusion window of 20 s was applied.

MeroX (Götze *et al*, 2015) was used to search the raw data. Searches were performed against an ad hoc protein database containing the sequences of GroEL, GroES, β-galactosidase and 70S ribosomal proteins as well as a set of randomized decoy sequences generated by MeroX with the minimum peptide length set to 5 amino acids, maximum number of missed cleavages set to 3 and False Discovery Rate cut-off set to 5%. Variable modifications were set to carbamidomethylation of cysteine (mass shift 57.02146 Da) and methionine oxidation (mass shift 15.99491 Da). DSBU modified fragments correspond to 85.05276 Da and 111.03203 Da (precision: 5 ppm MS and 10 ppm MS/MS). Crosslinks to non-native N-terminal sequences (present in purified RNCs because of N-terminal purification tag cleavage) and crosslinks with scores below 50 were disregarded. xiVIEW online tool (Graham *et al*, 2019) was used to visualise crosslinks on linear domain diagrams. Detected crosslinks are listed in Table S3.

### Negative stain electron microscopy

GroEL:RNC and EL:ES_2_:RNC complexes prepared via co-sedimentation assays described above were diluted to 12 nM in RNC low-salt buffer. 4 µL were applied to copper grids with 300 mesh carbon film (EM Resolutions) that had been glow discharged (25 mA, 60s) in the GloQube Plus Glow Discharge System (Quorum Technologies). After 1-2 minutes of incubation, excess sample was removed with filter paper and 2% uranyl acetate was applied three times for 20 s. Excess stain was removed with filter paper and grids were air-dried. Around ∼500 micrographs were collected for each dataset with a pixel size of 4.3 Å/pixel, and defocus range of -1 to -2.2 µm on a FEI Tecnai Spirit 120 kV TEM. Data-processing was performed using cryoSPARC (Punjani *et al*, 2017). Following CTF estimation, particles were automatically picked using the blob picker and classified in 2D. 2D classes corresponding to GroEL particles were used as templates for a second auto-picking. For each analysed dataset, between 30,000 and 40,000 GroEL or GroEL:ES_2_ particles were selected after at least two rounds of 2D classification. Particles were subsequently used to generate an initial 3D model. After at least two rounds of 3D classification, two 3D classes in each dataset were selected for 3D refinement. The number of particles in each class is stated in each figure. EM map docking, visualization and figures were prepared using UCSF ChimeraX (Pettersen *et al*, 2021).

## Data availability

The mass spectrometry proteomics data have been deposited to the ProteomeXchange Consortium via the PRIDE partner repository (Perez-Riverol *et al*, 2022) with the following dataset identifiers: Composition of ribosomal pellets: PXD054251 HDX-MS: PXD054376 XL-MS: PXD054379

## Acknowledgements

We thank F.U. Hartl (MPI Biochemistry) for the plasmids encoding TF, DnaK and DnaJ; John Christodoulou (University College London) for the *Δtig* E. coli strain; Svend Kjaer for preparing GFP-clamp resin; Grant Pellowe and Stephane Mouilleron (Francis Crick institute) for purified 3C and TEV proteases; Santosh Shivakumaraswamy for purified full-length β-gal; Tomas Voisin for assistance with the nsEM experiments; Christelle Soudy (Francis Crick Institute Chemical Biology STP), Grant Pellowe, Santosh Shivakumaraswamy and Karim El bouri for help preparing immobilised pepsin; Steven Howell for proteomic analysis of the ribosomal pellets, and all members of the Protein Biogenesis Lab (Francis Crick Institute) for help and discussion. This work is supported by the Francis Crick Institute which receives its core funding from Cancer Research UK (CC2025, CC1063, CC1065, CC1068), the UK Medical Research Council (CC2025, CC1063, CC1065, CC1068), and the Wellcome Trust (CC2025, CC1063, CC1065, CC1068). For the purpose of Open Access, the author has applied a CC BY public copyright licence to any Author Accepted Manuscript version arising from this submission.

## Author contributions

A.R. designed and performed most of the experiments, analysed the data and wrote the manuscript together with D.B. S.L.M. and J.M.S. collected and processed the XL-MS data. G.J., J.H. and R.E. purified GroEL and GroES. D.B. conceived and supervised the project.

## Supplementary Material

**Figure S1.**
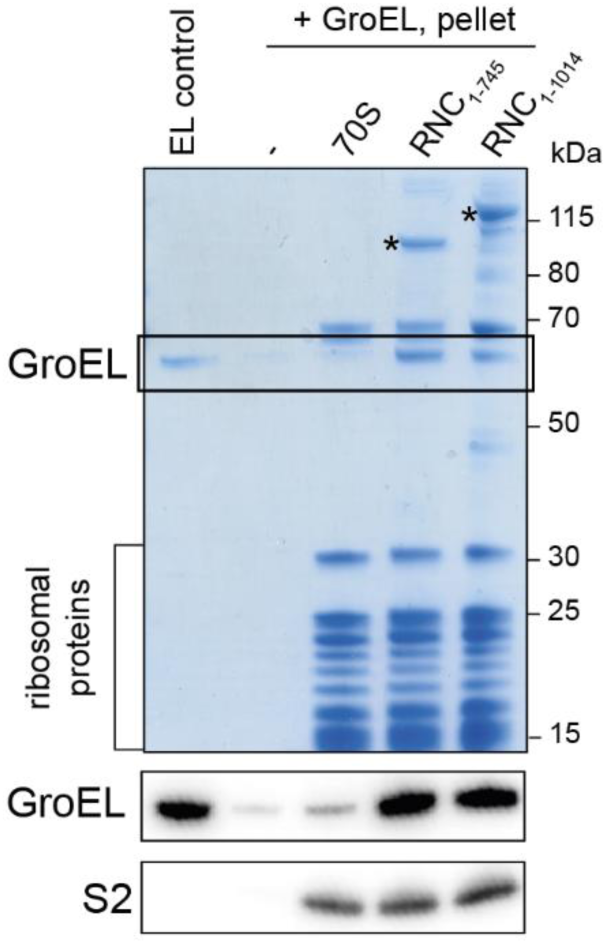
GroEL co-sediments with RNCs. Top: Coomassie-stained SDS-PAGE of the resuspended ribosomal pellet from a co-sedimentation assay. Prior to sedimentation, GroEL was incubated with either buffer (-), empty ribosomes (70S), RNC_1-745_, or RNC_1-1014_. RNCs were purified from *Δtig* cells. Bands corresponding to the NCs (*) and GroEL are indicated. Purified GroEL was loaded in the first lane for reference (EL control). Bottom: immunoblot of a replicate SDS-PAGE gel probed using antibodies against GroEL and ribosomal protein S2.

**Figure S2.**
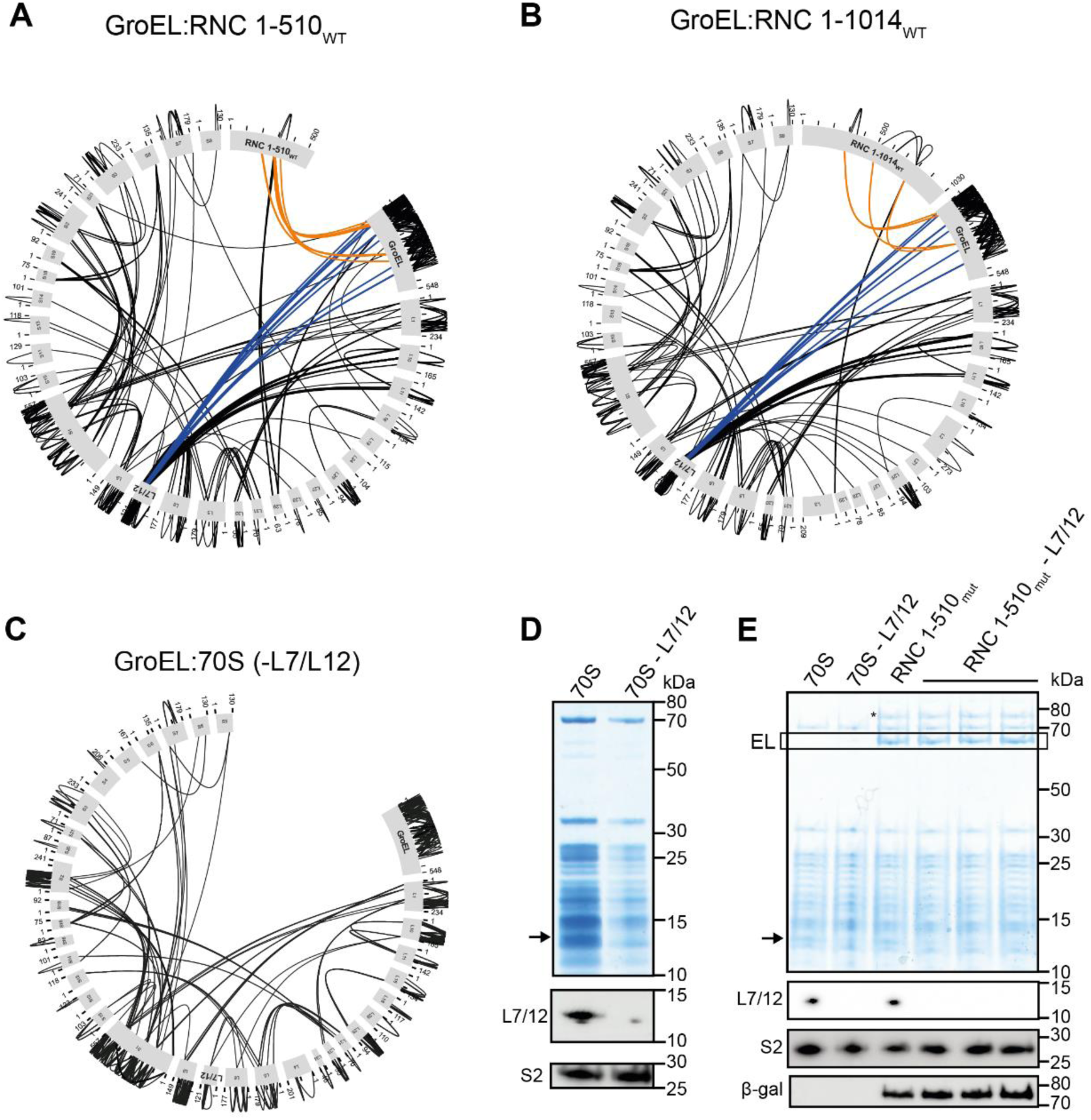
GroEL crosslinking to empty ribosomes and RNCs. **(A)** Map of crosslinks between GroEL and RNC1-510WT. Crosslinks between GroEL and the NC (orange) or L7/L12 (blue) are highlighted. **(B)** Map of crosslinks between GroEL and RNC_1-1014_. Crosslinks between GroEL and the NC (orange) or L7/L12 (blue) are highlighted. **(C)** Map of crosslinks between GroEL and empty 70S ribosomes depleted of L7/L12. **(D)** Selective removal of L7/L12 from ribosomes. Top: Coomassie-stained SDS-PAGE of complete (70S) and L7/L12-depleted (70S-L7/12) ribosomes. The position of L7/12 is indicated by a black arrow. Bottom: Immunoblot of a replicate SDS-PAGE gel probed using antibodies against ribosomal proteins L7/12 and S2. **(E)** Removing L7/L12 from RNC_1-510mut_ does not prevent GroEL binding. Top: Coomassie-stained SDS-PAGE of the resuspended ribosomal pellet from a co-sedimentation assay. Prior to sedimentation, GroEL was incubated with either empty ribosomes (70S), 70S ribosomes after depletion of the ribosomal stalk (70S-L7/12), RNC_1-510mut_ (RNC 1-510_mut_), or RNC 1-510_mut_ after depletion of the ribosomal stalk (RNC 1-510_mut_-L7/12). The pelleting assay for the last condition (RNC 1-510_mut_-L7/12) was conducted in triplicate. Bands corresponding to the NCs (*) and GroEL are indicated. Bottom: Immunoblot of a replicate SDS-PAGE gel probed using antibodies against β-galactosidase and ribosomal proteins L7/12 and S2.

**Figure S3.**
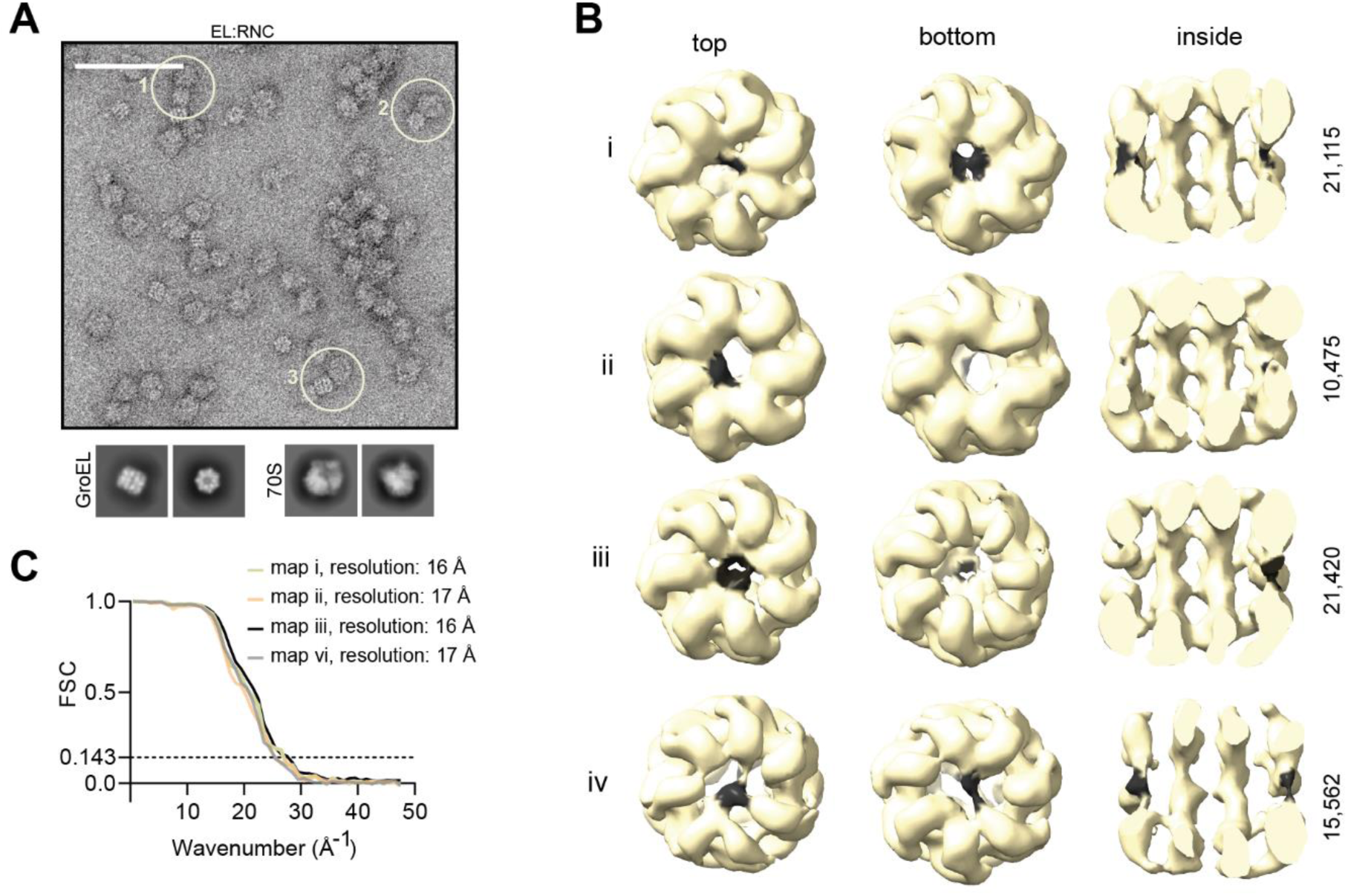
nsEM of GroEL:RNC_1-510_ complexes. **(A)** Top: Negative stain electron microscopy (nsEM) micrographs of uncrosslinked GroEL:RNC1-510 complexes. The scale bar corresponds to 100 nm. Examples of GroEL positioned near ribosomes are circled (1-3). Bottom: 2D class averages of GroEL and 70S ribosomes. **(B)** 3D reconstructions of uncrosslinked (i, ii) and DSBU-crosslinked (iii, iv) GroEL: RNC_1-510_ complexes from nsEM. Density not accounted for by the solved structure of GroEL (PDB: 5W0S) is coloured black. The number of particles contributing to each reconstruction is given on the right. **(C)** Fourier Shell Correlation (FSC) plots for reconstructions obtained from the uncrosslinked (maps i and ii) and DSBU-crosslinked (maps iii and iv) complexes.

**Figure S4.**
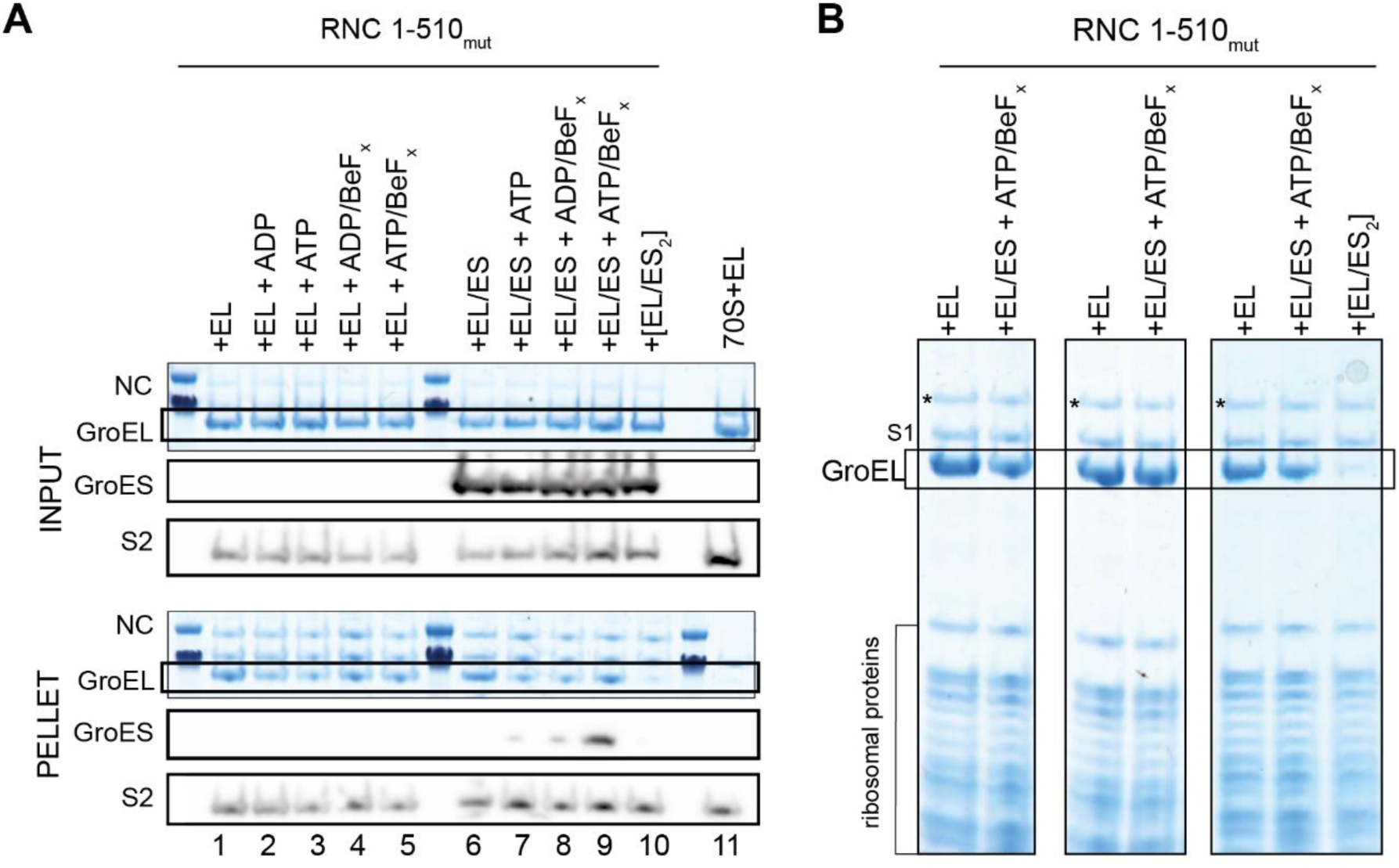
GroEL/ES binding to RNC_1-510mut_ in the presence of different nucleotides. **(A)** Effect of different nucleotides on the stability of GroEL:RNC complexes. Coomassie-stained SDS-PAGE and immunoblot analysis of co-sedimentation assays of GroEL/ES with RNC_1-510mut_ incubated with different nucleotides. Both input (top) and pellet (bottom) fractions are shown. Prior to sedimentation, the RNC was incubated with GroEL either in low-salt RNC buffer (1), or with additional 1 mM ADP (2), ATP (3), ADP/BeF_x_ (4) or ATP/BeF_x_ (5). Alternatively, the RNC was incubated with GroEL/ES in low-salt RNC buffer (6) or with additional 1 mM ATP (7), ADP/BeF_x_ (8) or ATP/BeF_x_ (9). As controls, the RNC was incubated with a pre-formed complex of EL:ES_2_ (10) in the presence of ATP/BeF_x_, or GroEL was incubated with empty 70S ribosomes (11). Any nucleotide and metal salts were present in the binding buffer as well as the sucrose cushion and wash buffers. Bands corresponding to the NCs (*) and GroEL are highlighted. Below each Coomassie-stained gel are immunoblots from the same gel probed using antibodies against GroES and ribosomal protein S2. **(B)** GroEL remains bound to RNCs upon addition of GroES and ATP/BeF_x_. Coomassie-stained SDS-PAGE of resuspended ribosomal pellets from co-sedimentation assays of GroEL with RNC_1-510mut_. Where indicated, RNCs were incubated with GroEL, or GroEL, GroES and ATP/BeF_x_. As a control, GroEL, GroES and ATP/BeF_x_ were pre-mixed to form symmetrically closed complexes before adding to RNCs (+[EL:ES]_2_). Bands corresponding to the NCs (*) and GroEL are highlighted. Each lane corresponds to an independent co-sedimentation assay.

**Figure S5.**
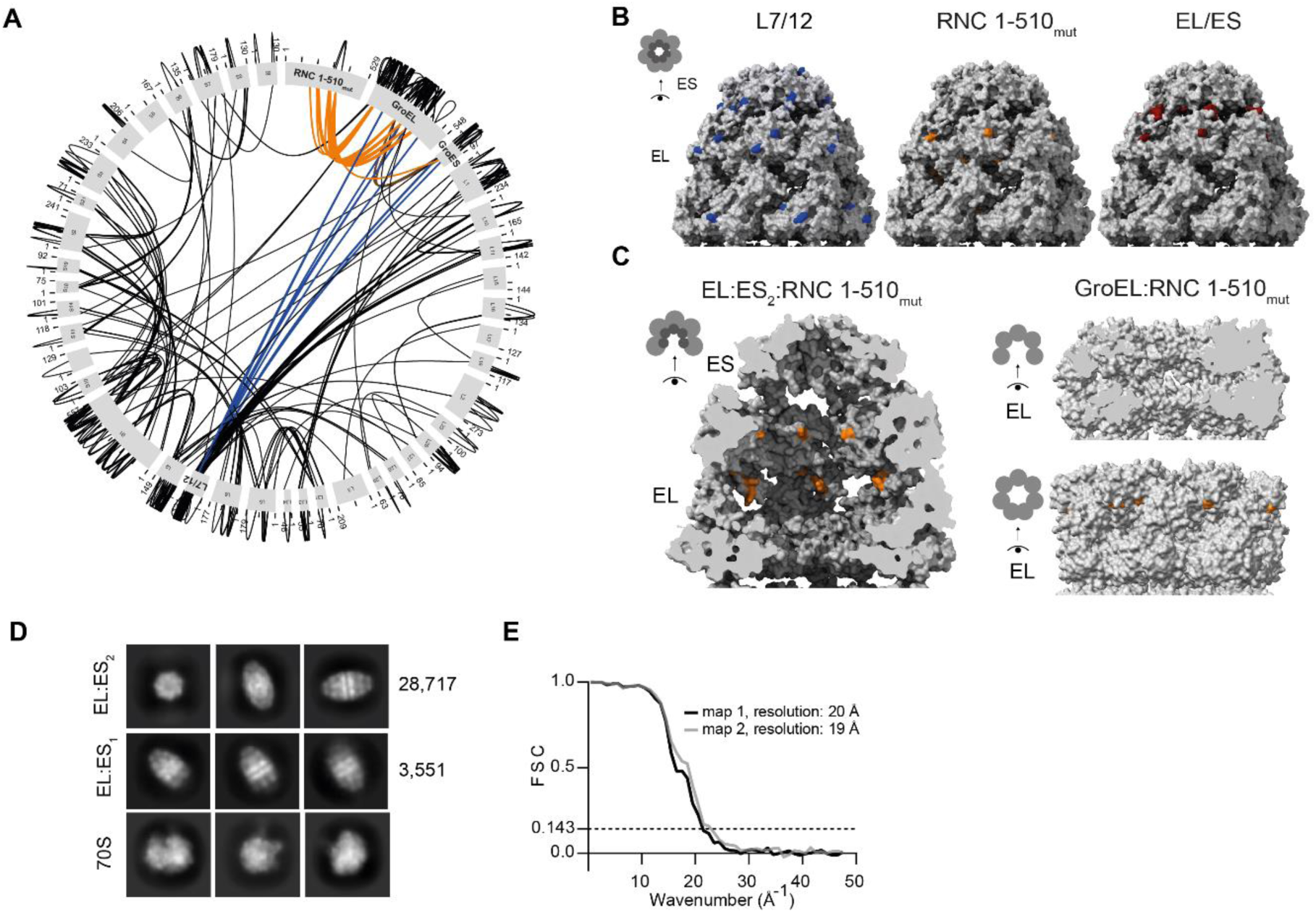
Structural characterisation of the GroEL:ES_2_:RNC complex. **(A)** Map of crosslinks between GroEL/ES and RNC1-510mut. Crosslinks between GroEL and GroES (brown), from GroEL/ES to the NC (orange), and from GroEL/ES to L7/L12 (blue) are highlighted. **(B)** Crosslink sites are mapped onto the structures of the ATP/BeFx-stabilised EL:ES2 complex (PDB:7VWX), showing the outer surface. Residues are separated according to whether they crosslink to L7/L12 (blue), the NC (orange), or connect GroEL and GroES (brown). **(C)** Change in accessibility of GroEL residues upon GroES binding. Left: inner surface of the GroEL/ES cavity (left, PDB:7VWX). Right: inner (top) and outer (bottom) surfaces of apo-GroEL (PDB: 5W0S). Residues which crosslinked to the NC in the EL:ES_2_:RNC complex but not in the GroEL:RNC complex are shown in orange. **(D)** 2D class averages for double-capped GroEL (EL:ES2), single-capped GroEL (EL:ES1) and 70S ribosomes, from nsEM of the GroEL/ES:RNC complex. **(E)** Fourier Shell Correlation (FSC) plots for reconstructions obtained from nsEM analysis of EL:ES2:RNC complexes.

**Figure S6.**
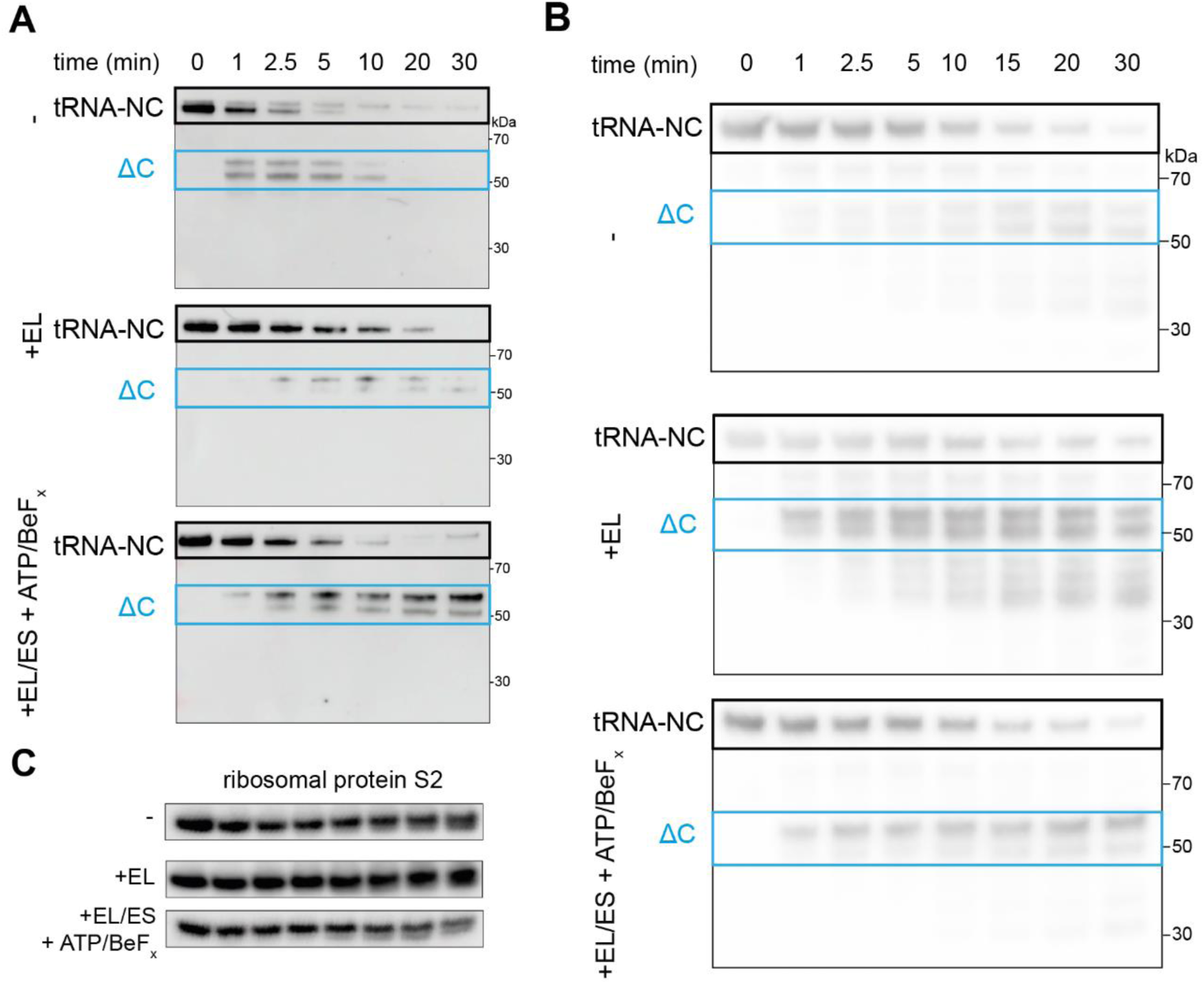
Limited proteolysis of RNC_1-510mut_. **(A)** Replicate of limited proteolysis experiment shown in Figure 6, using an antibody raised against the N-terminus of β-galactosidase. **(B)** As in Figure 6, except GroEL/ES:RNC complexes were isolated by sedimentation through a sucrose cushion, prior to treatment with proteinase K. **(C)** Loading control for limited proteolysis experiment shown in (B). Immunoblots were probed using an antibody against ribosomal protein S2.

**Figure S7.**
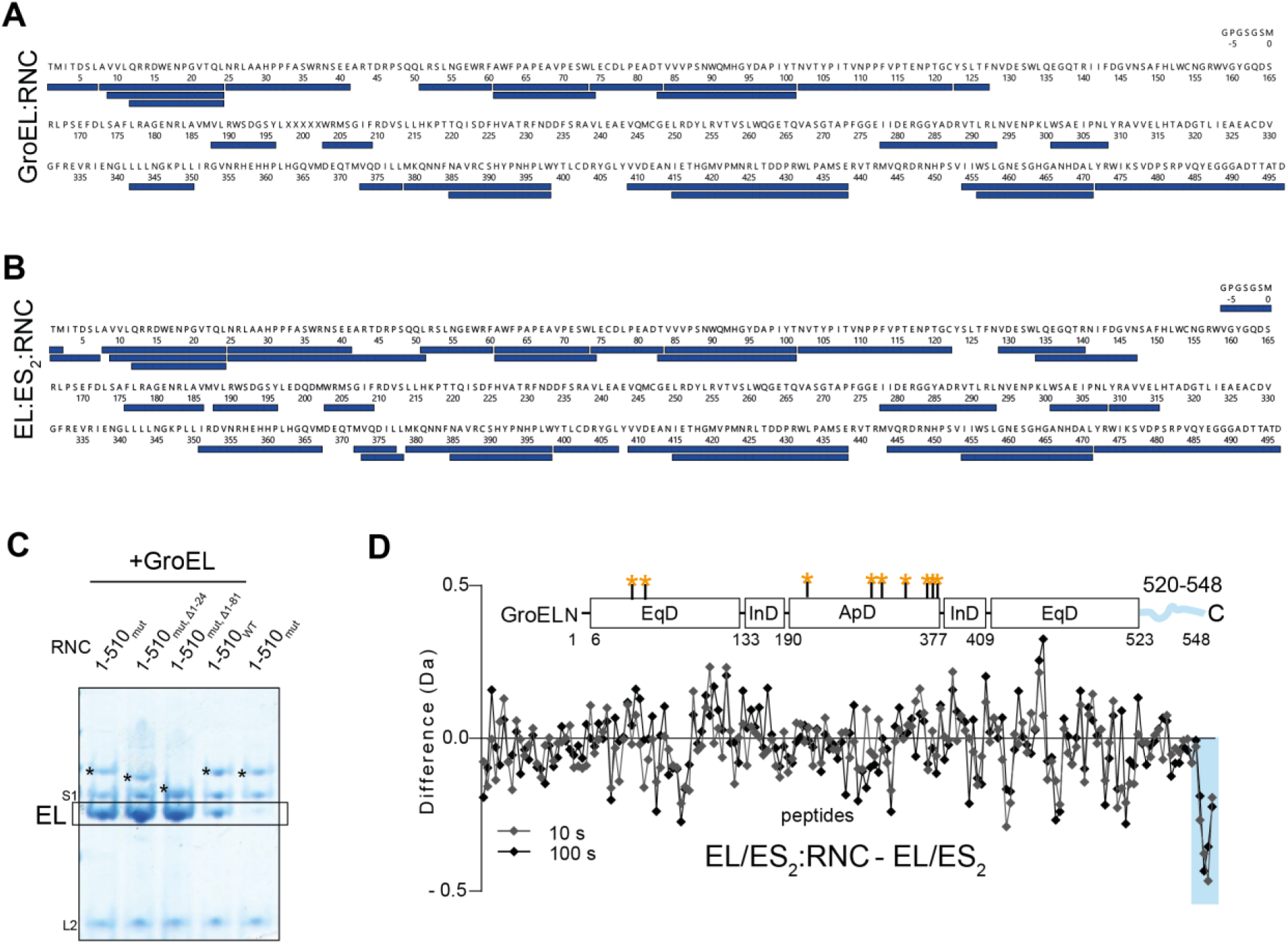
GroEL/ES binding to RNCs. **(A)** Peptide coverage map of the NC in GroEL:RNC1-510mut complex. **(B)** Peptide coverage map of the NC in EL:ES2 RNC1-510mut complex. **(C)** GroEL binding is unaffected by deleting residues 1-81 of NC_1-510mut._ Coomassie-stained SDS-PAGE of the resuspended ribosomal pellet from a co-sedimentation assay. Prior to sedimentation, GroEL was incubated with either wild-type RNC_1-510_ (1-510_WT_), RNC_1-510mut_ (1-510_mut_), or RNC_1-510mut_ lacking residues 1-24 (1-510_mut Δ1-24_) or 1-81 (1-510_mut Δ1-81_). The final lane contains purified RNC 1-510_mut_ without additional GroEL. Bands corresponding to the NCs (*) and GroEL are indicated. **(D)** Protection of GroEL upon binding GroES RNC1-510mut. Difference in deuterium uptake after 10 s (grey) or 100 s (black), between isolated EL:ES2 and EL:ES2:RNC1-510mut, both stabilised by ATP-BeFx. Values are plotted for individual GroEL peptides. Negative values indicate less deuteration of a peptide in when the RNC is bound.

**Figure S8.**
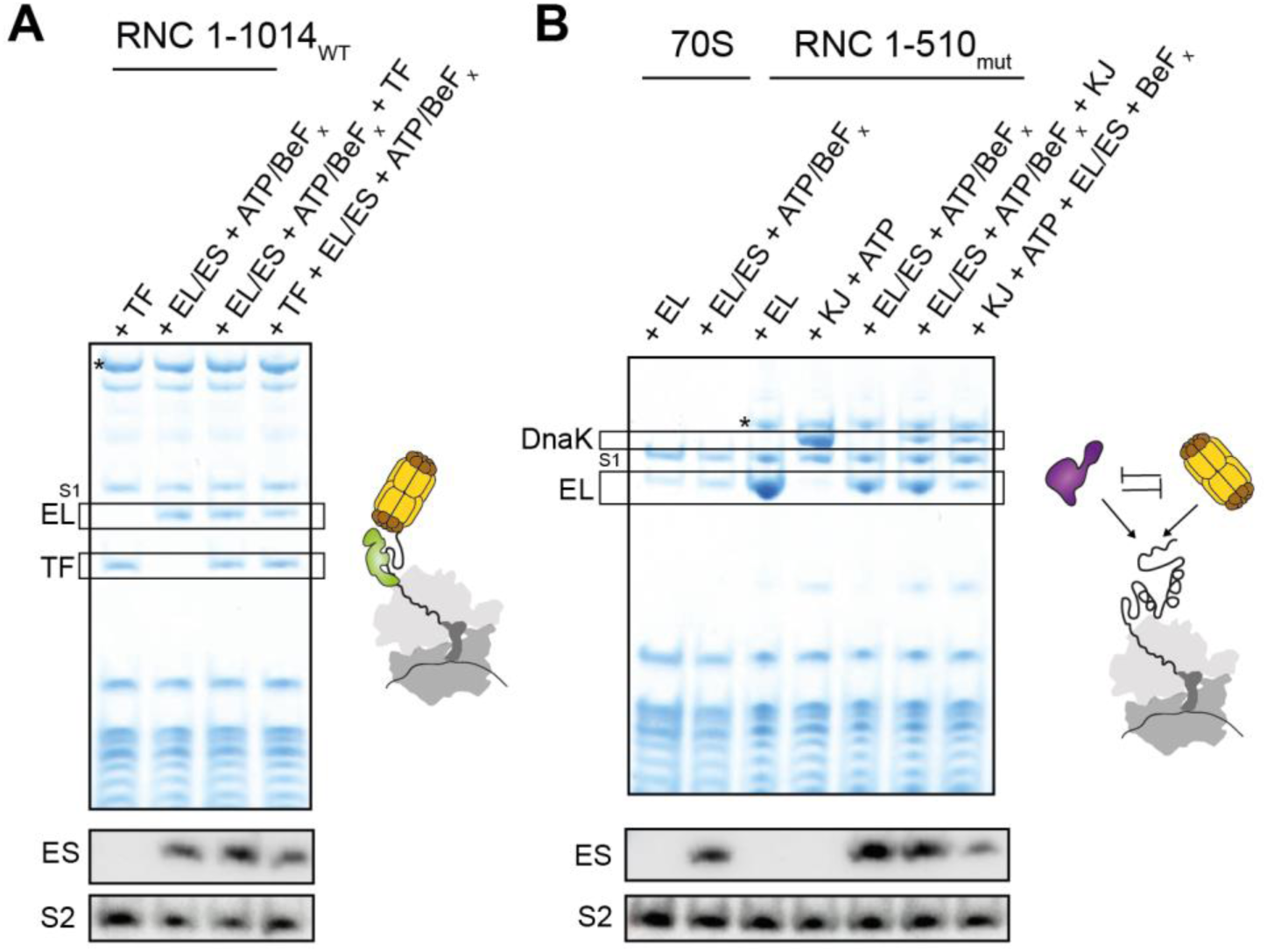
Coordination of GroEL/ES with Trigger factor and DnaK. **(A)** TF and GroEL/ES do not compete for binding long NCs. Top: Coomassie-stained SDS-PAGE of the resuspended ribosomal pellet from co-sedimentation assays. Prior to sedimentation, RNC1-1014 was incubated with Trigger factor (+TF), GroEL with GroES and ATP/BeFx (+EL/ES), or a combination of the above in the specified order. Bands corresponding to the NC (*), TF and GroEL are indicated. Bottom: Immunoblot from an equivalent SDS-PAGE gel, probed against GroES and ribosomal protein S2. **(B)** DnaK and GroEL/ES compete for binding NCs. Top: Coomassie-stained SDS-PAGE of the resuspended ribosomal pellet from co-sedimentation assays. Prior to sedimentation, empty ribosomes (70S) or RNC1-510mut were incubated with GroEL (+EL), DnaK with DnaJ (+KJ), GroEL with GroES and ATP/BeFx (+EL/ES), or a combination of the above in the specified order. Bands corresponding to the NC (*), DnaK and GroEL are indicated. Bottom: Immunoblot from an equivalent SDS-PAGE gel, probed against GroES and ribosomal protein S2.

## Supplementary table legends

**Table S1** HDX-MS data

**Table S2** Proteomic analysis of resuspended pellets from co-sedimentation assays

**Table S3** XL-MS data

**Table S4** List of DNA constructs and recombinant protein sequences

